# The homeobox transcription factor DUXBL controls exit from totipotency

**DOI:** 10.1101/2022.09.19.508541

**Authors:** Maria Vega-Sendino, Teresa Olbrich, Paula Stein, Desiree Tillo, Grace I. Carey, Virginia Savy, Bechara Saykali, Catherine N. Domingo, Tapan K. Maity, Lisa M. Jenkins, Carmen J. Williams, Sergio Ruiz

## Abstract

Upon exit from the totipotent 2-cell (2C) embryo stage, the 2C-associated transcriptional program needs to be efficiently silenced. However, the molecular mechanisms involved in this process remain mostly unknown. Here, we demonstrate that the 2C-specific transcription factor DUX directly induces the expression of DUXBL to promote this silencing. Indeed, DUX expression in *Duxbl*-knockout ESC causes increased induction of the 2C-transcriptional program, whereas DUXBL overexpression impairs 2C-associated transcription. CUT&RUN analyses show that DUXBL gains accessibility to DUX-bound regions in DUX-induced ESC while it is unable to bind those regions in uninduced cells. Mechanistically, we determined that DUXBL interacts with TRIM24 and TRIM33, two members of the tripartite motif superfamily involved in gene silencing and co-localizes with them in nuclear foci upon DUX expression. Furthermore, DUXBL downregulation in mouse zygotes leads to a penetrant 2C-stage arrest. Our data reveals an unexpected role for DUXBL in controlling the exit from totipotency.

## INTRODUCTION

Totipotency, the ability of a single cell to develop into a full organism, is a property strictly associated with the zygote and the 2C embryo in mice^1^. This unlimited potential becomes restricted upon further development and zygotic genome activation (ZGA) seems to be the trigger for loss of totipotency *in vivo*^2-4^. ZGA is the process that determines the transition from maternal to zygotic transcription. In mice, it starts as a minor wave from S-phase of the zygote to G1 phase of the 2C embryo and is followed by a major transcriptional wave during the mid-to-late 2C embryo stage^2-4^. Interestingly, ZGA confers to the 2C embryo a unique transcriptional profile characterized by the expression of cleavage stage-specific genes, including *Dux*, genes from the *Zscan4(a-f)* cluster, *Zfp352*, and *Pramef*-family genes, and the murine endogenous retroviral element (MERVL)-family of transposable elements (TE)^5-7^. These TE become derepressed following the wave of global demethylation after fertilization and are highly transcribed at the 2C stage. Their long-terminal repeats (LTR) act as functional promoters, generating long chimeric transcripts with both retroviral and endogenous gene sequences including putative open reading frames^8^. Although the functional relevance of these sequences is still a matter of debate, interfering with their expression leads to developmental defects^9^. Despite the quick activation of 2C-associated genes and MERVL elements during ZGA, their silencing is required upon the exit from totipotency at the 4C stage to proceed with development. Indeed, sustained activation of the 2C-associated transcriptional program induces developmental arrest with blastomeres at the 4C stage having a 2C transcriptional signature^10^. Silencing of the 2C transcriptional program involves redundant but rather independent mechanisms, including transcriptional repression of the *Dux* gene by TRIM66, the E3 ligase PIAS4 or SMCHD1 as well as physical recruitment of *Dux* loci to nucleolar heterochromatin^11-14^. Furthermore, MERVL repeats are transcriptionally silenced by the protein complex ZMYM2/LSD1 or the chromatin remodeler CAF-1 and post-transcriptionally by the RNA decay complex NEXT^15-17^. However, whether a direct DUX-dependent negative feedback loop mechanism ensures a timed and irreversible silencing of the 2C-associated transcriptional program remains unknown.

The protein DUX is a pioneering transcription factor that is necessary and sufficient to trigger a ZGA-associated transcriptional signature in embryonic stem cells (ESC)^5-7^. Indeed, expression of DUX increases the natural occurrence of 2C-like cells (2CLC), endogenous cells within ESC cultures with transcriptional and chromatin features associated with 2C embryos^5-7^. Interestingly, the human ortholog DUX4 is involved in the facioscapulohumeral muscular dystrophy, a disorder characterized by elevated levels of DUX4 and reactivation of endogenous retroviral sequences and human ZGA transcripts in myotubes leading to pronounced cell toxicity^18^. Similarly, sustained DUX expression in mouse cells induces cell death and extensive DNA damage^19^. Surprisingly, despite the prominent role of DUX in triggering a ZGA-associated signature *in vitro, Dux*-deficient mice are viable with mild pre- and post-implantation defects^20,21^. These observations revealed that DUX is important but largely dispensable for ZGA in mice and suggested that additional transcription factors, such as OBOX4, could compensate for its loss^22^.

DUX belongs to the family of double homeobox domain transcription factors and thus, it contains two N-terminal DNA-binding homeodomains in addition to a C-terminal sequence that confers transcriptional activation through the recruitment of the histone acetyl transferases (HAT) CBP/p300^23^. Indeed, deletion of this C-terminal region (DUX*Δ*C) abolishes DUX-mediated gene transactivation and cell toxicity suggesting that this interaction is necessary to mediate the DUX-induced transcriptional program^23^. Another member of the DUX family is DUXBL, located in a near-telomeric cluster on chromosome 14 and only found in the mouse and rat genomes^24,25^. DUXBL lacks the C-terminal domain observed in DUX and conserved in human DUX4, suggesting that DUXBL does not interact with CBP/p300 and, similar to DUX*Δ*C lacks transcriptional activation potential^24, 25^. DUXBL is involved in β-selection by inducing apoptosis in pre-T cells with failed rearrangements^26^. Moreover, DUXBL over-expression in muscle cells is sufficient to initiate tumorigenesis by promoting mesenchymal-to-epithelial transition. In fact, the *Duxbl* gene is amplified in p53-deficient rhabdomyosarcoma^27^. Nevertheless, the role of DUXBL during early development remains unknown.

In this work, we show that DUXBL is transcriptionally induced by DUX and is efficiently recruited together with the TRIM24/TRIM33 complex to DUX-enriched regions to promote transcriptional silencing and heterochromatinization. Indeed, DUXBL overexpression in ESC leads to impaired 2C-like conversion while *Duxbl*-knockout ESC show an increased 2C-associated transcriptional response. Importantly, DUXBL knockdown in mouse embryos leads to a complete arrest at the 2C-stage. Our results reveal an unexpected role for DUXBL in controlling the exit from totipotency.

## RESULTS

### DUX induces two protein forms from the *Duxbl* gene

We previously generated ESC carrying a doxycycline (DOX)-inducible *Dux* cDNA (hereafter, ESC^DUX^) and used this model to examine *Duxbl* induction^19^. We observed that *Dux* expression induced *Duxbl* in a dose-dependent manner (Fig. 1a). Interestingly, *Duxbl* follows the wave of *Dux* expression during early embryonic development (Fig. 1b)^28^. Using published DUX-HA ChIP-seq datasets^5^ we confirmed that DUX binds three regions upstream the start codon of the *Duxbl* gene (Fig. 1c). To explore whether these binding regions (BR1, 2 and 3) contribute to *Duxbl* gene expression, we generated fluorescence-based reporter vectors by subcloning each BR to control the expression of a GFP gene copy in a PiggyBAC construct. In addition, we examined MERVL reactivation using an LTR-RFP reporter system in which RFP expression is driven by a specific LTR from MERVL sequences^8^. LTR-RFP reporter ESC^DUX^ were then stably transfected with the corresponding PiggyBAC construct to determine GFP expression upon DOX treatment. We observed that BR1 was constitutively active in ESC, independent of DUX expression (Extended Data Fig. 1a). However, BR2 and BR3 showed DUX-dependent activation with expression levels of GFP correlating with those of RFP (Fig. 1d and Extended Data Fig.1b). We next investigated the activation dynamics of these BR during the 2CLC conversion process. We observed that BR2 and BR3 become active after DUX expression but before full conversion to a 2CLC-state (Supplementary Videos 1 and 2).

**Fig. 1:**
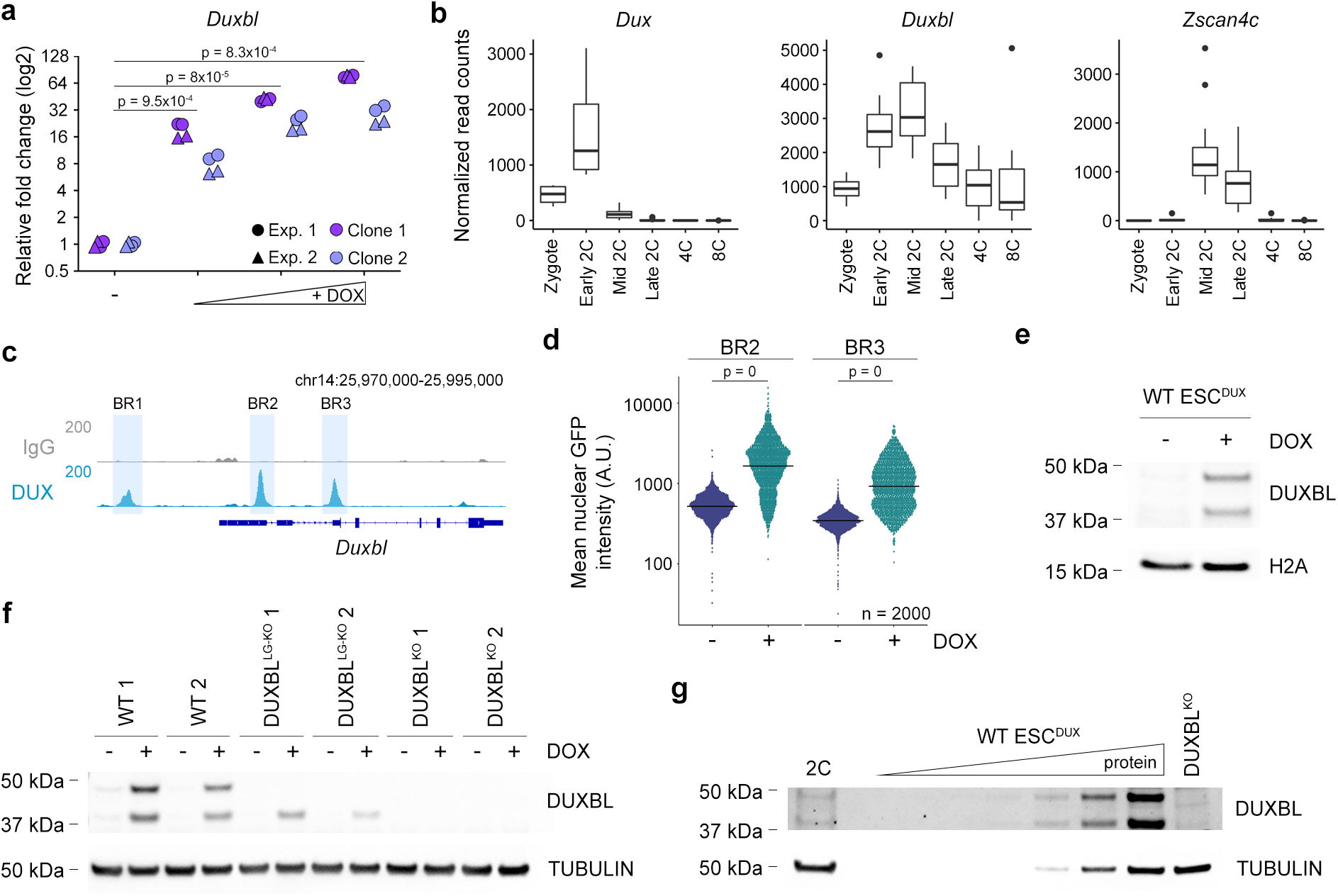
The *Duxbl* gene generates two protein products upon DUX expression. **a)** Relative fold change (log2) expression of *Duxbl* upon DUX expression in ESC^DUX^. Reactions were performed in duplicate with two independent experiments using two different ESC^DUX^ lines. p values are shown from two-tailed paired *t*-tests. **b)** Plots showing normalized read counts for *Duxbl, Dux* and *Zscan4c* during early development. *Zscan4c* was included as an example of a DUX-dependent gene during ZGA. Single cell RNAseq data was extracted from^28^. Center line indicates the median, box extends from the 25^th^ to 75^th^ percentiles and whiskers extend from the hinge to the largest or smallest value no further than 1.5-fold from the inter-quartile range. Data beyond the whiskers are considered outliers and plotted individually. **c)** Genome browser tracks showing DUX occupancy at the *Duxbl* gene. ChIP-seq data obtained from^5^. DUX binding sites and BR regions are highlighted. Read mapping was performed using the mm39 mouse genome version. **d)** High-throughput imaging quantification of GFP mean nuclear intensity in untreated or DOX-treated for 16 hours BR2 or BR3-GFP reporter ESC^DUX^. Center lines indicate mean values. n=2000; p values are shown from one-tailed unpaired *t*-tests. **e)** Western blot analysis of DUXBL performed in lysates from untreated or DOX-treated for 16 hours ESC^DUX^. H2A levels are shown as a loading control. **f)** Western blot analysis of DUXBL performed in lysates from untreated or DOX-treated for 16 hours WT, DUXBL^LG-KO^ and DUXBL^KO^ ESC^DUX^. Tubulin levels are shown as a loading control. **g)** Western blot analysis of DUXBL performed in lysates from a pool of approximately one thousand late 2C-embryos. Increasing amounts of DOX-treated WT ESC^DUX^ as well as DOX-treated DUXBL^KO^ ESC^DUX^ lysates were included as controls. Tubulin levels are shown as a loading control. For (e-g) two independent experiments were performed but one representative experiment is shown.

To examine DUXBL expression, we generated a custom rabbit polyclonal antibody against the C-terminal region of DUXBL. Although DUXBL was not expressed in uninduced ESC, we were surprised that our Western blot analysis consistently showed the presence of two bands of equal intensity in DUX-induced ESC (Fig. 1e). The upper band corresponds to a protein form with the predicted molecular weight for DUXBL (DUXBL large form, DUXBL^LG^) while the lower band could represent a shorter DUXBL protein form. Indeed, mass spectrometry analyses confirmed the identity of the protein of lower molecular weight as DUXBL (DUXBL small form, DUXBL^SM^; Supplementary Table 1). We noticed an ATG at the end of the second exon that could act as an alternative start codon generating a putative protein form with the expected molecular weight of DUXBL^SM^ following alternative splicing. Therefore, to investigate the origin of DUXBL^SM^, we performed a rapid amplification of cDNA ends (RACE) analysis to examine the different RNA molecules originating from the *Duxbl* gene after DUX expression. Although we detected two major RNA molecules originating from BR2 and BR3, they both contained the first exon (Extended Data Fig. 1c). Still, RNA molecules with a non-canonical Kozak sequence, which is the case for *Duxbl*, could randomly use an alternative start codon downstream of the predicted ATG to generate different protein forms^29^. To confirm this possibility, we used Cas9-based editing with a sgRNA targeting the second exon in *Duxbl* with the aim of inducing frameshift mutations upstream of the second ATG or eliminating the alternative start codon. In fact, we determined that small deletions upstream of the second ATG led to the unique expression of DUXBL^SM^ (DUXBL^LG-KO^ ESC^DUX^; Fig. 1f and Extended Data Fig. 1d). However, larger deletions affecting the proper splicing of exon 2 and/or eliminating the alternative ATG led to a full *Duxbl* knockout in ESC (DUXBL^KO^ ESC^Dux^; Fig. 1f and Extended Data Fig. 1d). Interestingly, the resulting DUXBL^SM^ lacked most of the first homeodomain (Extended Data Fig. 1e). These data demonstrated that the ATG in exon 2 is a functional start codon and both DUXBL^LG^ and DUXBL^SM^ originate from the same RNA molecule. Finally, we examined whether DUXBL was expressed in mouse embryos. Unfortunately, our custom antibody was not useful to detect endogenous levels of DUXBL in embryos by immunofluorescence. However, Western blot analyses of late 2C embryos following the wave of DUX expression revealed expression of both DUXBL^LG^ and DUXBL^SM^, confirming that embryos express the two DUXBL protein products (Fig. 1g).

### DUXBL^KO^ ESC show increased activation of the 2C transcriptional program

To determine the role of DUXBL in 2CLC conversion, we first examined the dynamics of endogenous 2CLC conversion in wild type (WT) and DUXBL^KO^ ESC. Interestingly, the residency time in the 2CLC state after conversion was considerably extended in DUXBL^KO^ ESC (Fig. 2a and Extended Data Fig. 2a). Moreover, DUXBL deficiency promoted a slight increase in endogenous 2CLC conversion (Extended Data Fig. 2b). We next performed RNA-seq in untreated or DOX-treated WT ESC^DUX^, DUXBL^LG-KO^ ESC^DUX^ or DUXBL^KO^ ESC^DUX^. Consistent with previous reports^5-7^, DUX induced the 2C-associated transcriptional program in all ESC lines (Extended Data Fig. 2c). To examine the consequences of DUXBL loss, we focused our analysis on comparing DOX-treated samples as DUXBL would only be expressed in that condition. We only observed a few genes differentially expressed in DOX-treated DUXBL^LG-KO^ ESC^DUX^ compared to WT (Fig. 2b and Supplementary Table 2). Similarly, we did not detect expression changes in most TE families (Fig. 2b and Supplementary Table 2). These data suggest that the expression of DUXBL^SM^ might be sufficient to compensate for the loss of DUXBL^LG^. Indeed, we identified a few hundred genes differentially expressed in DOX-treated DUXBL^KO^ ESC^DUX^ compared to WT (Fig. 2c and Supplementary Table 2). Moreover, MERVL elements were also significantly upregulated in DOX-treated DUXBL^KO^ ESC^DUX^ (Fig. 2c and Supplementary Table 2). Interestingly, the expression of the set of genes upregulated in DOX-treated DUXBL^KO^ ESC^DUX^ was enriched in the 2C embryo compared to other developmental stages while the expression of the downregulated genes did not show enrichment at any specific stage (Fig. 2d, Extended Data Fig. 2d). Furthermore, these upregulated genes showed a significant overlap with genes induced by DUX expression in ESC (Fig. 2e,f). Altogether, these results demonstrate that DUXBL^KO^ ESC^DUX^ show an exacerbated induction of 2C-associated genes and TE upon DUX expression suggesting that DUXBL limits the activation of the DUX-induced transcriptional program.

**Fig. 2:**
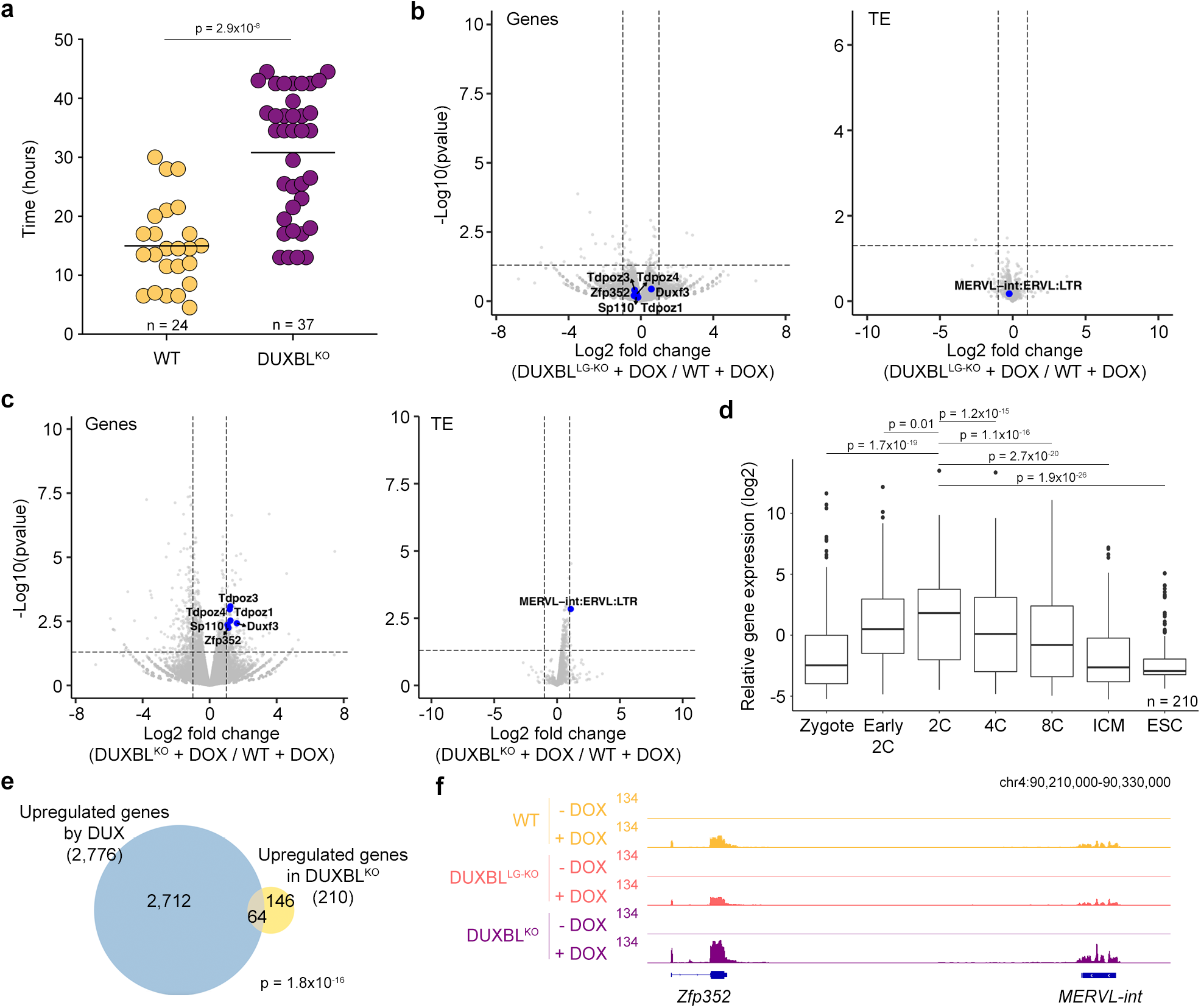
DUXBL deficiency leads to increased activation of the 2C-associated program. **a)** Graph showing the time that DOX-treated WT and DUXBL^KO^ ESC^Dux^ stay in a 2CL state evaluated by using the LTR-RFP reporter. p value is shown from a two-tailed unpaired *t*-test. **b, c)** Volcano plots showing differentially expressed genes and TE between DOX-treated DUXBL^LG-KO^ (b) or DUXBL^KO^ ESC^DUX^ (c) compared to DOX-treated WT ESC^DUX^. Relevant genes and TE are highlighted. **d)** Box and whisker plot showing normalized fold change (log2) expression of the 210 genes upregulated in DOX-treated DUXBL^KO^ ESC^DUX^ compared to DOX-treated WT ESC^DUX^ during preimplantation development including ESC. RNAseq data obtained from^36^. p values are shown from two-tailed unpaired *t*-tests. Center line indicates the median, box extends from the 25^th^ to 75^th^ percentiles and whiskers extend from the hinge to the largest or smallest value no further than 1.5-fold from the inter-quartile range. Data beyond the whiskers are considered outliers and plotted individually. **e)** Venn diagram showing the overlap between the 210 genes described in (c) and the upregulated genes in LTR^+^ sorted DUX-expressing ESC (data obtained from^5^). p value was obtained from a hypergeometric test. **f)** Genome browser tracks showing RNAseq RPKM read count at the indicated region in untreated or DOX-treated WT, DUXBL^LG-KO^ and DUXBL^KO^ ESC^DUX^.

### The expression level of DUXBL determines 2CLC conversion efficiency

To further assess the role of DUXBL in limiting 2CLC conversion we generated DOX-inducible WT ESC lines expressing DUXBL^LG^ (Extended Data Fig. 3a). We then tested the influence of DUXBL expression by inducing 2CLC reprogramming using the spliceosome inhibitor pladienolide B (PlaB) or retinoic acid (RA) added to the culture media^30,31^. Interestingly, we observed that 2CLC conversion mediated by PlaB or RA treatment was further increased in DUXBL^KO^ ESC compared to WT ESC (Fig. 3a, b). Conversely, expression of DUXBL^LG^ reduces the percentage of 2CLC in both PlaB and RA-treated cells (Fig. 3c, d). Consistently, Western blot and real-time PCR analysis showed a significant decrease in the expression of 2C-associated genes upon induction of DUXBL^LG^ in PlaB-or RA-treated WT ESC (Fig. 3e, f). We also generated DOX-inducible DUXBL^LG^ as well as DUXBL^KO^ cell lines in an auxin-inducible degron line for *Ctcf* (ESC^CTCF-AID^) (Extended Data Fig. 3a,b) as CTCF depletion has been shown to efficiently induce 2CLC conversion^19^. Expression of DUXBL^LG^ impaired 2CLC conversion whereas an increase in the percentage of RFP^+^ cells was observed in DUXBL^KO^ ESC^CTCF-AID^ after CTCF depletion (Extended Data Fig. 3c,d). DUX interacts with and recruits the HATs CBP/P300, which are necessary for proper activation of the 2C transcriptional program^23^. Thus, we examined the levels of H3K27ac in PlaB-treated cells and observed a decrease in the levels of H3K27ac at DUX-bound regions upon DUXBL^LG^ expression (Fig. 3g). Our results demonstrate that DUXBL expression limits 2CLC conversion.

**Fig. 3:**
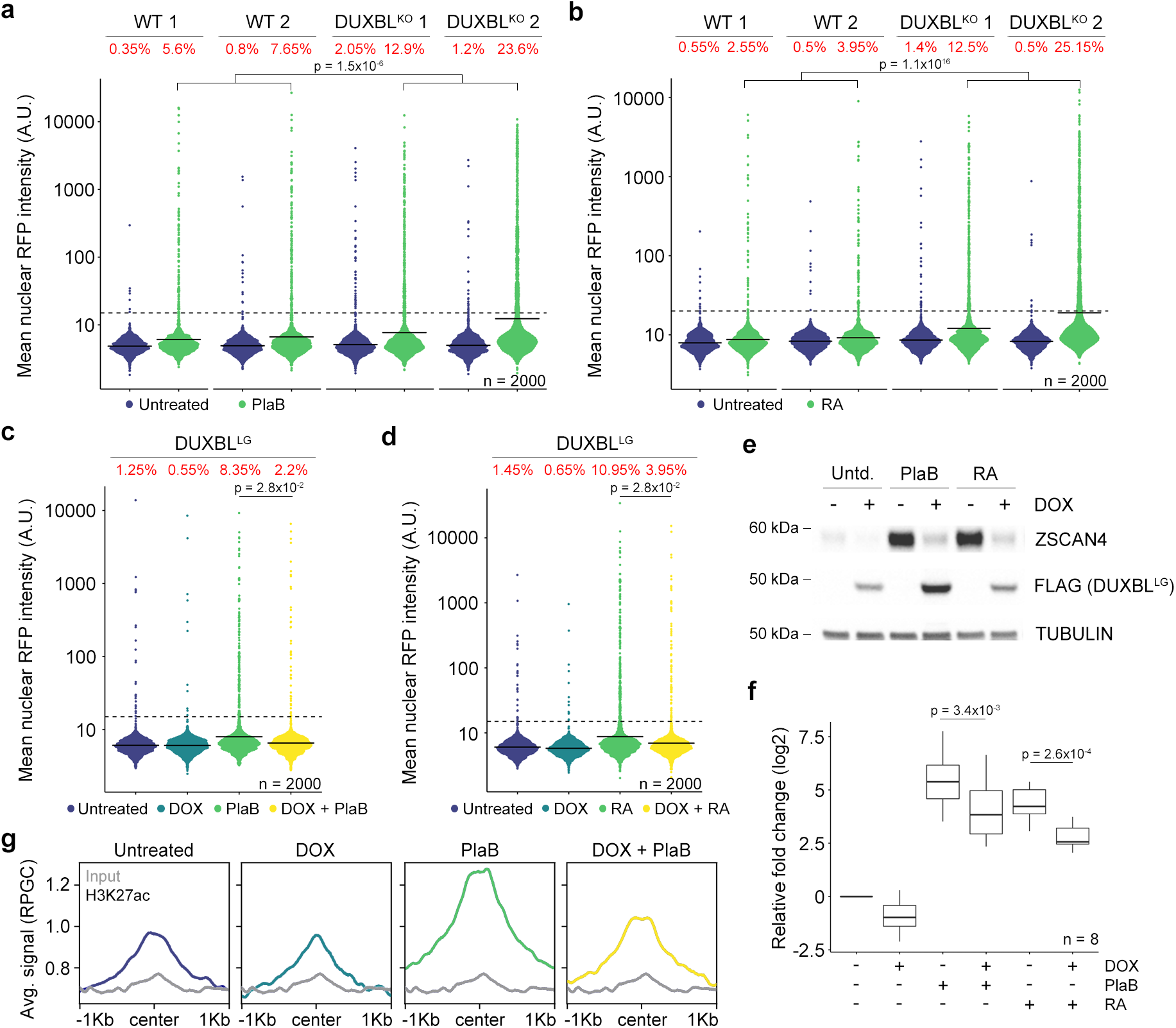
2CLC conversion efficiency is affected by DUXBL levels. **a, b)** High-throughput imaging quantification of RFP^+^ cells in LTR-RFP reporter WT and DUXBL^KO^ incubated with 2.5 µM PlaB for 24 hours (a) or 0.53 µM RA for 48 hours (b). Center lines indicate mean values. Percentages of RFP^+^ cells above the threshold (dotted line) are indicated. n=2000; p value is shown from one-tailed unpaired *t*-test. Three independent experiments were performed but only one representative is shown. **c, d)** High-throughput imaging quantification of RFP^+^ cells in untreated or DOX-treated LTR-RFP reporter ESC^DUXBL-LG^ incubated with 2.5 µM PlaB for 24 hours (c) or 0.53 µM RA for 48 hours (d). Center lines indicate mean values. Percentages of RFP^+^ cells above the threshold (dotted line) are indicated. n=2000; p value is shown from one-tailed unpaired *t*-test. Three independent experiments were performed but only one representative is shown. **e)** Western blot analysis of the indicated proteins performed using lysates from untreated or DOX-treated ESC^DUXBL-LG^ incubated with 2.5 µM PlaB for 24 hours or 0.53 µM RA for 48 hours. Two independent experiments were performed but only one representative is shown. **f)** Box and whisker plot showing the relative fold change (log2) expression of eight 2C-associated genes (*Dux, Zscan4c, Zfp352, Tcstv3, Sp110, Tdpoz1, Dub1* and *Eif1ad8*) in untreated or DOX-treated ESC^DUXBL-LG^ incubated with 2.5 µM PlaB for 24 hours or 0.53 µM RA for 48 hours. Two independent experiments were performed but only one representative is shown. p values are shown from one-tailed unpaired *t*-tests. Center line indicates the median, box extends from the 25^th^ to 75^th^ percentiles and whiskers show Min to Max values. **g)** CUT&RUN read density plot (reads per genome content, RPGC) showing H3K27ac enrichment at DUX-bound sites in untreated or DOX-treated WT ESC expressing DUXBL^LG^ incubated with 2.5 µM PlaB for 24 hours. Input (IgG) is shown as reference control.

### DUXBL gains chromatin accessibility to DUX-bound regions upon DUX expression

DUX has been proposed to be a pioneering transcription factor based on its ability to remodel and promote chromatin opening, including at 2C-associated genes and MERVL elements, when expressed in ESC^5^. Thus, we next sought to examine the DUXBL binding landscape in ESC by native CUT&RUN sequencing using an antibody against DUXBL. For this purpose, we established ESC lines constitutively expressing DUXBL^LG^. We identified a total of 2353 overlapping peaks between two independent DUXBL^LG^-expressing ESC lines (Supplementary Table 3). These peaks were significantly enriched for the predicted binding sequence for DUXBL (Extended Data Fig. 4a and Supplementary Table 4)^32^. Furthermore, we captured over 70% of these peaks by using a FLAG antibody on DOX-inducible ESC expressing a FLAG-tagged version of DUXBL^LG^ (ESC^DUXBL-LG^; Extended Data Fig. 4b,c and Supplementary Table 5). DUXBL^LG^ bound the promoters of hundreds of genes (Fig. 4a-c). Interestingly, DUXBL^LG^ binding was particularly enriched at ESC enhancers (507/2353 DUXBL^LG^ peaks) and super-enhancers (76/2353 DUXBL^LG^ peaks) (Fig. 4a,d,e)^33,34^. Gene ontology analysis of DUXBL^LG^-bound regions showed significant enrichment for processes such as “chromatin organization and assembly”, “cellular response to leukemia inhibitory factor” and “blastocyst development” (Fig. 4f). We next examined the chromatin binding overlap between DUXBL and DUX as they bind a similar DNA sequence (Extended Data Fig. 4a). Interestingly, we found little overlap between DUX and DUXBL^LG^ binding sites (317/2353 DUXBL^LG^ peaks) and no enrichment at MERVL elements (Fig. 4g and Extended Data Fig. 4d). Indeed, DUXBL^LG^ preferentially bound to genomic regions having an open chromatin configuration compared to DUX, which bound mostly inaccessible regions in ESC as determined by ATAC-seq analysis (Extended Data Fig. 4e,f)^35^.

**Fig. 4:**
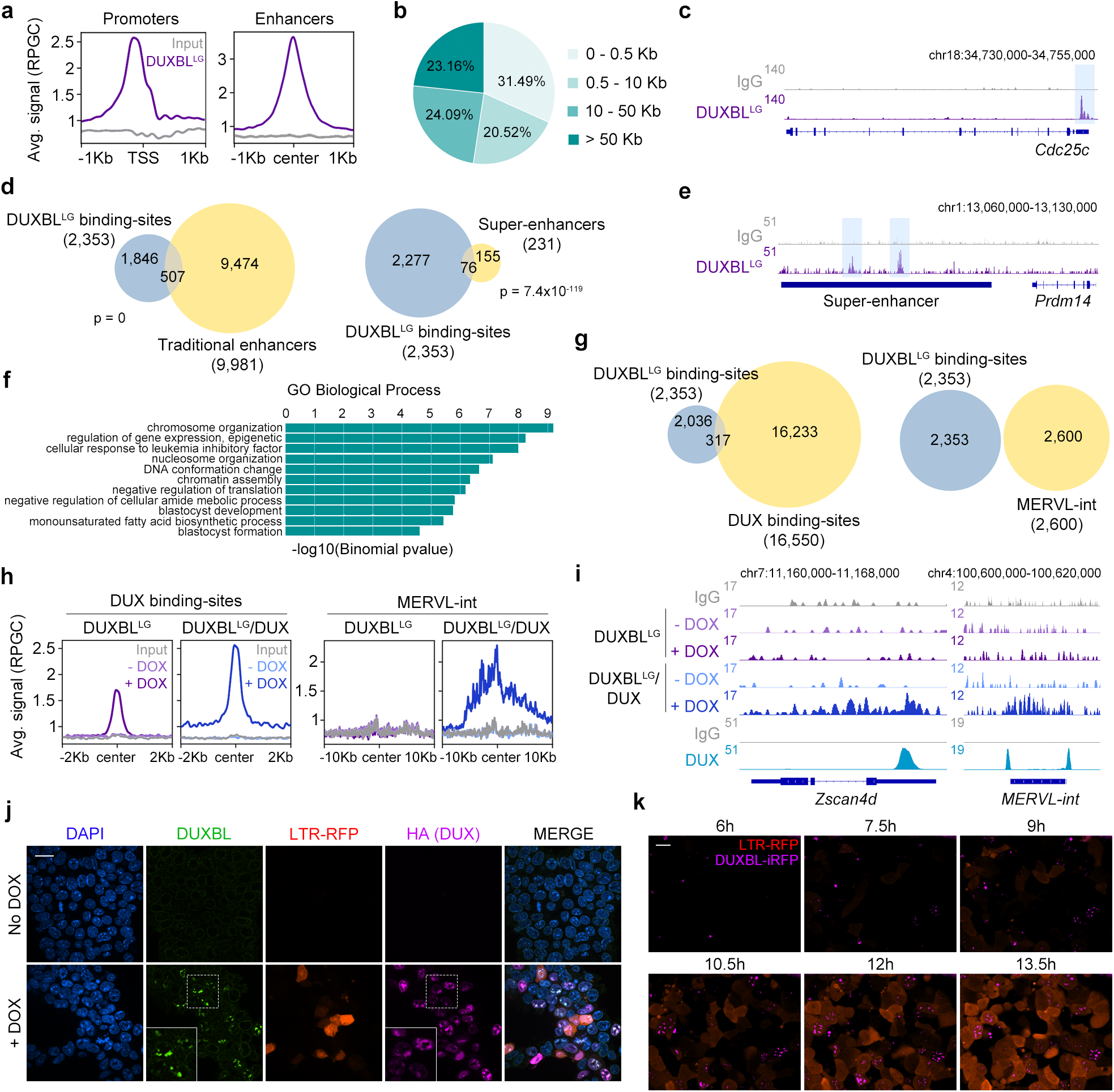
DUX-mediated chromatin opening facilitates DUXBL binding to DUX-bound regions. **a)** CUT&RUN normalized read density plot (reads per genomic content, RPGC) showing DUXBL^LG^ enrichment at gene transcriptional start sites (TSS, left panel) and traditional ESC enhancers (right panel) in WT ESC expressing DUXBL^LG^. Input (IgG) is shown as reference control. **b)** Pie chart showing the genomic distance of DUXBL^LG^ peaks from gene TSS. **c)** Genome browser tracks corresponding to samples from (a) showing DUXBL^LG^ occupancy at the indicated genomic region with a representative example of DUXBL^LG^ binding to a gene promoter. DUXBL^LG^ binding site is highlighted. Input (IgG) is shown as reference control. **d)** Venn diagrams showing the number of DUXBL^LG^ peaks overlapping with traditional ESC enhancers (left) and ESC super-enhancers (SE, right). Enhancer data obtained from^33,34^. p value shown is from Fisher’s exact test. **e)** Genome browser tracks corresponding to samples from (a) showing DUXBL^LG^ occupancy at the indicated genomic region with a representative example of DUXBL^LG^ binding to a SE. DUXBL^LG^ binding site and the SE are highlighted. Input (IgG) is shown as reference control. **f)** Gene ontology (GO) terms obtained from analyzing DUXBL^LG^ peaks. **g)** Venn diagrams showing the number of DUXBL^LG^ peaks overlapping with DUX-binding sites (left) and MERVL-int repeats (right). **h)** CUT&RUN read density plot (RPGC) showing DUXBL^LG^ enrichment at the indicated genomic regions in DOX-treated WT ESC expressing DUXBL^LG^ or DUX/DUXBL^LG^ as shown. Input (IgG) is shown as reference control. **i)** Genome browser tracks corresponding to samples from (h) showing DUX (data from^5^) and DUXBL^LG^ enrichment (our data) at the indicated genomic region with a representative example of DUXBL^LG^ binding to *Zscan4d* and a MERVL-int repeat. Input (IgG) is shown as reference control. **j)** Immunofluorescence analysis of DUX (HA) and endogenous DUXBL in LTR-RFP reporter DOX-treated ESC^DUX^. DAPI was used to visualize nuclei. Scale bar, 20 μm. Dashed regions are zoomed-in in the insets. **k)** Time lapse microscopy experiment performed in LTR-RFP reporter DOX-induced ESC^DUX^ endogenously tagged with a miRFP702 fluorescent protein at the *Duxbl* locus. Time since the addition of DOX is indicated. Scale bar, 20 μm.

Based on the sequential expression of DUX followed by DUXBL during ZGA and the sequence similarities between their binding sites, we hypothesized that DUX expression, and the subsequent chromatin opening in 2C-associated regions, could facilitate DUXBL binding to an expanded subset of binding sites. Thus, we generated DOX-inducible ESC lines expressing DUXBL^LG^ or DUXBL^LG^/DUX and found that DUX-bound regions showed increased accessibility to DUXBL^LG^ following DUX expression (Fig. 4h,i). Furthermore, DUXBL^LG^ was particularly enriched at MERVL sequences in DUX-expressing ESC (810/2600 MERVL-int elements) (Fig. 4h,i). When we examined the subcellular localization of endogenous DUXBL in DOX-induced ESC^DUX^, we found DUXBL co-localized with DUX in nuclear foci (Fig. 4j). Similar localization to nuclear foci was observed for exogenous DUXBL^LG^ in DUX-expressing ESC (Extended Data Fig. 4g). Importantly, DUXBL translocation to nuclear foci was dependent on the expression of DUX as we observed pan-nuclear distribution of DUXBL in DUXBL^LG^-expressing ESC in the absence of DUX (Extended Data Fig. 4h). We next examined the translocation dynamics of endogenous DUXBL by tagging the *Duxbl* locus with a fluorescent protein (iRFP702) in LTR-RFP reporter ESC^DUX^. Upon DUX expression, we observed DUXBL induction and nuclear foci formation early in the 2CLC conversion process, prior to full MERVL expression (Fig. 4k and Supplementary Video 3). These data combined show that DUXBL is efficiently translocated to chromatin regions specifically enriched for DUX binding.

### DUXBL interacts with TRIM33 and TRIM24

We next sought to investigate the mechanism by which DUXBL regulates the 2C-associated transcriptional program. We first used immunoprecipitation-mass spectrometry (IP-MS) analysis to examine the protein interactome of DUXBL in untreated and DOX-treated ESC^DUXBL-LG^. We detected a total of 160 proteins enriched in the DUXBL pulldown (fold change>5 as compared with the control, p value<0.05) including DUXBL (Fig. 5a, Extended Data Fig. 5a and Supplementary Table 6). However, we speculated that DUXBL interactors might differ when DUXBL is expressed in a 2CLC environment. Therefore, we also performed IP-MS analyses by pulling down endogenous DUXBL (DUXBL^LG^ and DUXBL^SM^) from untreated and DOX-treated ESC^DUX^ as well as only endogenous DUXBL (DUXBL^SM^) from untreated and DOX-treated DUXBL^LG_KO^ ESC^DUX^ and detected a total of 13 and 37 significantly enriched proteins, respectively (Extended Data Fig. 5b-e and Supplementary Table 6). Importantly, we identified specific interactors for each condition, suggesting context-specific protein interactions. Moreover, the existence of 26 unique interactors for DUXBL^SM^ revealed that DUXBL^LG^ and DUXBL^SM^ might have unique functions (Supplementary Table 6). Nevertheless, we observed a consistent interaction of DUXBL with the proteins TRIM24 and TRIM33 throughout all experimental conditions. TRIM24 and TRIM33 are related members of the transcriptional intermediary factor 1 of chromatin binding proteins and part of the larger tripartite-motif (TRIM) family of E3 ligases^37^. They both contain an N-terminal RING domain conferring E3 ligase activity and a plant homeodomain finger (PHD) and a bromodomain, which read methylation and acetylation status respectively on histone H3^37^. Moreover, TRIM24 contains a heterochromatin protein 1 (HP1) binding domain and interacts with histone deacetylases (HDAC)^37,38^. We first confirmed the interaction of DUXBL with TRIM24 and TRIM33 by IP-Western blot analysis using untreated and DOX-treated ESC^DUXBL-LG^ (Fig. 5b). In addition, although TRIM24 and TRIM33 showed pan-nuclear staining in untreated ESC, they translocated to nuclear foci upon DUX expression and co-localized with DUXBL and DUX (Fig. 5c,d and Extended Data Fig. 6a). Furthermore, the localization of TRIM24 in nuclear foci could also be observed in endogenously converted 2CLC (Extended Data Fig. 6b).

**Fig. 5:**
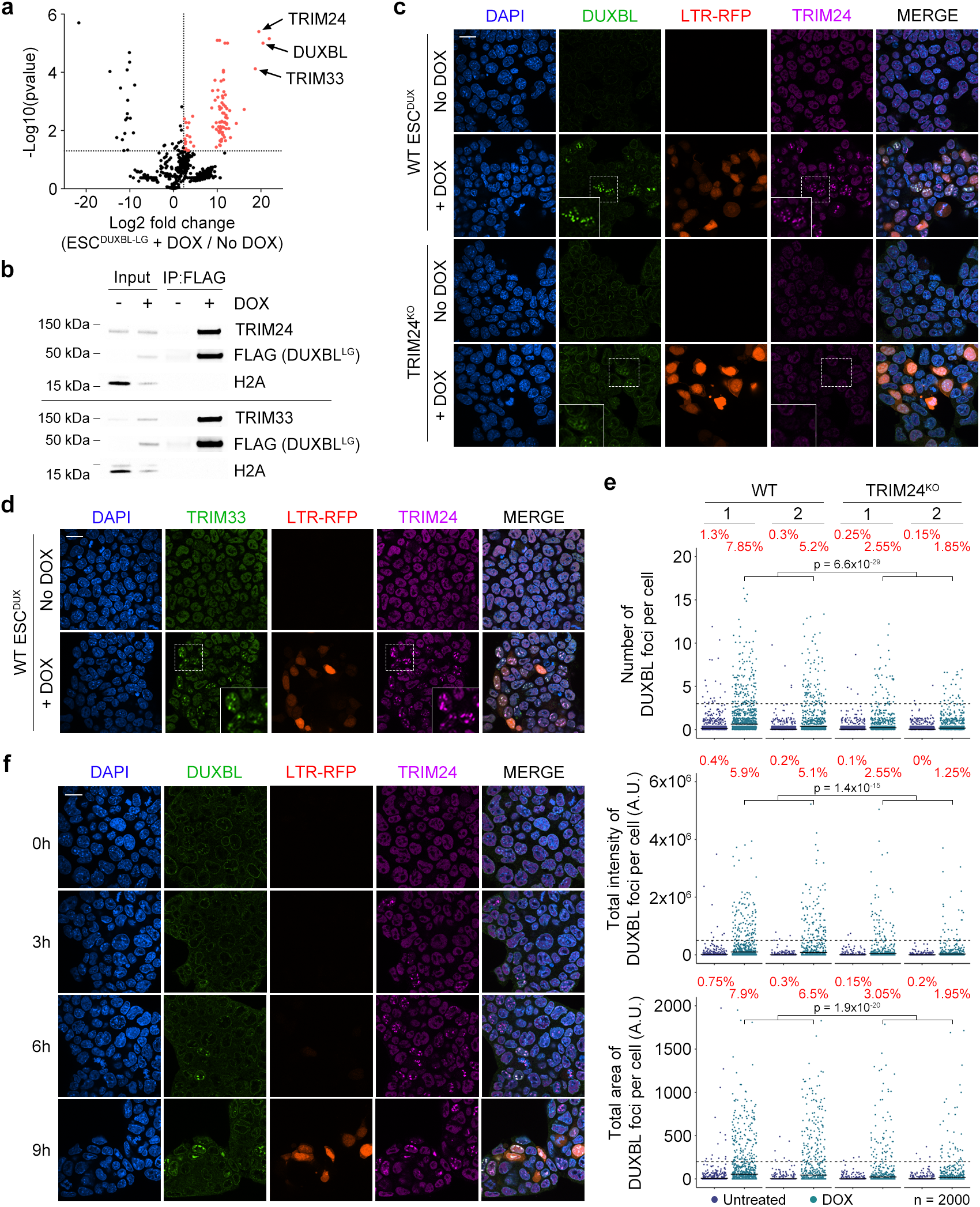
DUXBL interacts with the TRIM33/TRIM24 complex. **a)** Volcano plot showing the enrichment of proteins immunoprecipitated from DOX-treated over untreated ESC^DUXBL-LG^ by using two independent ESC lines from a representative DUXBL IP-MS experiment. Enriched proteins in DOX-treated cells are highlighted in red. **b)** Western blot analysis of the indicated proteins performed with FLAG immunoprecipitates obtained from untreated or DOX-treated ESC^DUXBL-LG^. **c)** Immunofluorescence analysis of endogenous DUXBL and TRIM24 in untreated or DOX-treated LTR-RFP reporter WT or TRIM24^KO^ ESC^DUX^. DAPI was used to visualize nuclei. Scale bars, 20 μm. Dashed regions are zoomed-in in the insets. Three independent experiments using two WT and TRIM24^KO^ ESC^DUX^ clones were performed but one representative experiment is shown. **d)** Immunofluorescence analysis of endogenous TRIM33 and TRIM24 in untreated or DOX-treated LTR-RFP reporter ESC^DUX^. DAPI was used to visualize nuclei. Scale bars, 20 μm. Dashed regions are zoomed-in in the insets. Three independent experiments using two ESC^DUX^ clones were performed but one representative experiment is shown. **e)** High-throughput imaging quantification of the number of DUXBL foci per cell (upper panel), the total intensity of DUXBL foci per cell (middle panel) and the total area of DUXBL foci per cell (lower panel) in untreated or DOX-treated LTR-RFP reporter WT or TRIM24^KO^ ESC^DUX^. Center lines indicate mean values. n=2000; Percentages above the threshold (dotted line) are indicated. Relevant p values are shown from one-tailed unpaired *t*-tests. Three independent experiments using at least two WT or TRIM24^KO^ ESC^DUX^ clones were performed but one representative experiment is shown. **f)** Immunofluorescence analysis of endogenous DUXBL and TRIM24 at different timepoints in untreated or DOX-treated WT ESC^DUX^. Time since the addition of DOX is indicated. DAPI was used to visualize nuclei. Scale bar, 20 μm.

We next asked whether DUXBL was necessary to promote TRIM24/33 recruitment. Thus, we examined the formation of TRIM24 nuclear foci in untreated and DOX-treated DUXBL^KO^ ESC^DUX^ and observed no differences in the number of foci compared to WT ESC (Extended Data Fig. 6c). We then generated TRIM24^KO^ and TRIM33^KO^ ESC^DUX^ (Extended Data Fig. 6d,e) and determined their ability to generate DUXBL nuclear foci upon DUX expression. We observed a significant reduction in DUXBL recruitment to foci in *Trim24*-knockout ESC^DUX^ but not in *Trim33*-knockout ESC^DUX^ suggesting that TRIM24 facilitates the recruitment of DUXBL to nuclear foci (Fig. 5c,e and Extended Data Fig. 6f). This can be observed by a decrease in the number of DUXBL nuclear foci as well as by a lower total DUXBL intensity at those foci (Fig. 5c,e). Furthermore, in agreement with a relevant role for TRIM24 in DUXBL recruitment, we observed lower levels of TRIM33 on a DUXBL-pulldown from DOX-induced *Trim24*-knockout ESC^DUX^ compared to control ESC^DUX^ (Extended Data Fig. 6g). When we examined the temporal dynamics of nuclear foci formation in DOX-induced ESC^DUX^, we observed that TRIM24 is quickly recruited within a few hours after DUX induction and is followed by DUXBL (Fig. 5f). Combined, these results show that DUXBL is recruited along with the repressor complex TRIM24/TRIM33 to DUX-bound regions.

### DUXBL/TRIM24/TRIM33 cooperate in the silencing of the 2C-transcriptional program

TRIM24 and TRIM33 physically interact with each other and are found complexed with the histone deacetylases HDAC1 and HDAC2, HP1 proteins and, to a lesser extent, to TRIM28^37,38^. Indeed, the TRIM24/33 protein complex is involved in the epigenetic silencing of retroviral elements in ESC^39,40^. Furthermore, ESC deficient in HP1 or TRIM28 or treated with HDAC inhibitors show increased 2CLC conversion^41-43^. We first examined whether HDAC1 and HP1 proteins were also enriched on TRIM24/33/DUXBL nuclear foci. Indeed, HDAC1 and HP1 were enriched at these foci suggesting they are also recruited upon DUX expression (Fig. 6a-d). The localization of HDAC and HP1 proteins on TRIM24/33/DUXBL nuclear foci suggested these regions might be undergoing H3K9me3 deposition and heterochromatin formation. Indeed, DUXBL-enriched chromatin regions identified in DUX-induced ESC^DUX^ showed a sharp H3K9me3 enrichment *in vivo* during the 2C to 4C embryo transition, especially at MERVL elements (Extended Data Fig. 7a,b)^44^. Therefore, we examined H3K9me3 deposition on DUX-induced nuclear foci and observed an increased accumulation (Fig. 6e). Furthermore, H3K9me3 deposition accumulated over time and correlated with TRIM24 recruitment after DUX expression (Fig. 6f and Extended Data Fig. 7c). These data suggested that DUX-induced nuclear foci undergo a progressive heterochromatinization as a response to the chromatin opening and transcriptional activation of genes and MERVL repeats induced by DUX.

We next asked whether the presence of DUX at the nuclear foci was required to maintain these foci. Thus, we generated DOX-inducible ESC lines expressing a FKBP-tagged DUX protein that can be degraded using dTAG compounds (ESC^DUX-FKBP^)^45^. DUX-FKBP protein induced 2CLC conversion in DOX-induced ESC^DUX-FKBP^ and treatment with dTAG compounds induced efficient degradation of the chimeric DUX protein (Fig. 6g and Extended Data Fig. 7d). To examine whether DUX degradation leads to nuclear foci dissolution, we analyzed DUXBL and TRIM24 retention at nuclear foci in untreated or dTAG-treated DOX-induced cells (Fig. 6g,h and Extended Data Fig. 7e). DUX-expressing cells showed consistent enrichment of DUX/DUXBL/TRIM24 at nuclear foci on LTR-RFP positive ESC. However, dTAG treatment of DOX-induced cells led to the dissolution of the nuclear foci with the consequent release of DUXBL and TRIM24 (Fig. 6g,h and Extended Data Fig. 7d,e). Importantly, LTR-RFP positive ESC showing residual DUX expression after dTAG treatment retained DUXBL on foci (Fig. 6g). Overall, these results demonstrate that DUX is necessary and sufficient to initiate the formation and maintenance of the nuclear foci, and its loss dictates their dissolution.

### DUXBL is required for embryos to exit from the 2C stage

Finally, we examined the role of DUXBL in mouse embryos *in vivo* by microinjecting morpholino oligonucleotides (MO) targeting the expression of both DUXBL^LG^ and DUXBL^SM^ in zygotes and followed embryonic development *in vitro*. By microinjecting MO in zygotes, we minimized the possibility of DUXBL maternal contribution in our experiments. We observed a highly penetrant 2C-arrest in *Duxbl*-knockdown embryos compared to non-injected or control MO-injected embryos (Fig. 7a,b). Similar results were obtained by using a second MO that targeted a distinct region, as well as by microinjecting siRNAs against *Duxbl* in oocytes followed by *in vitro* maturation and fertilization and assessment of embryo development (Extended data Fig. 8a,b). These siRNAs against *Duxbl* were highly efficient targeting *Duxbl* in DOX-treated ESC^DUX^ (Extended data Fig. 8c). These observations revealed that DUXBL is required to exit the 2C stage in mouse embryos. We next asked whether ZGA was taking place in *Duxbl*-knockdown embryos. Thus, we collected control or *Duxbl* MO-injected 2C embryos and performed RNA-seq. Principal component analysis (PCA) plot of these samples together with an available dataset of samples that evaluate temporal transcriptional dynamics during early mouse development^28^ showed that both control and *Duxbl* MO-injected embryos clustered with late 2C embryos (Fig. 7c). This confirms that DUXBL was not necessary to trigger ZGA and that *Duxbl*-knockdown embryos expressed embryonic transcripts as well as MERVL repeats. We next examined differentially expressed genes between control and *Duxbl* morpholino-injected embryos and observed 425 upregulated and 405 downregulated genes (Fold Change>2; p value<0.01, Fig. 7d and Supplementary Table 7). Interestingly, decreased DUXBL expression also led to upregulation of different classes of TE (Fig. 7e, Extended data Fig. 8d and Supplementary Table 7). Indeed, we observed that *Duxbl* MO-injected embryos showed a significantly increased percentage of the total amount of reads mapping to TE (Fig. 7f). Among the different classes of TE contributing to this increase, the SINE family was the largest contributor from an averaged 7% of the total reads to over 15% of the total reads in control or *Duxbl*-MO-injected embryos respectively (Fig. 7g). In particular, the TE family that showed the highest differential expression within the SINE class was the B2 family (Extended data Fig. 8e). In summary, DUXBL downregulation leads to a 2C-embryo arrest characterized by the upregulation of TE.

**Fig. 6:**
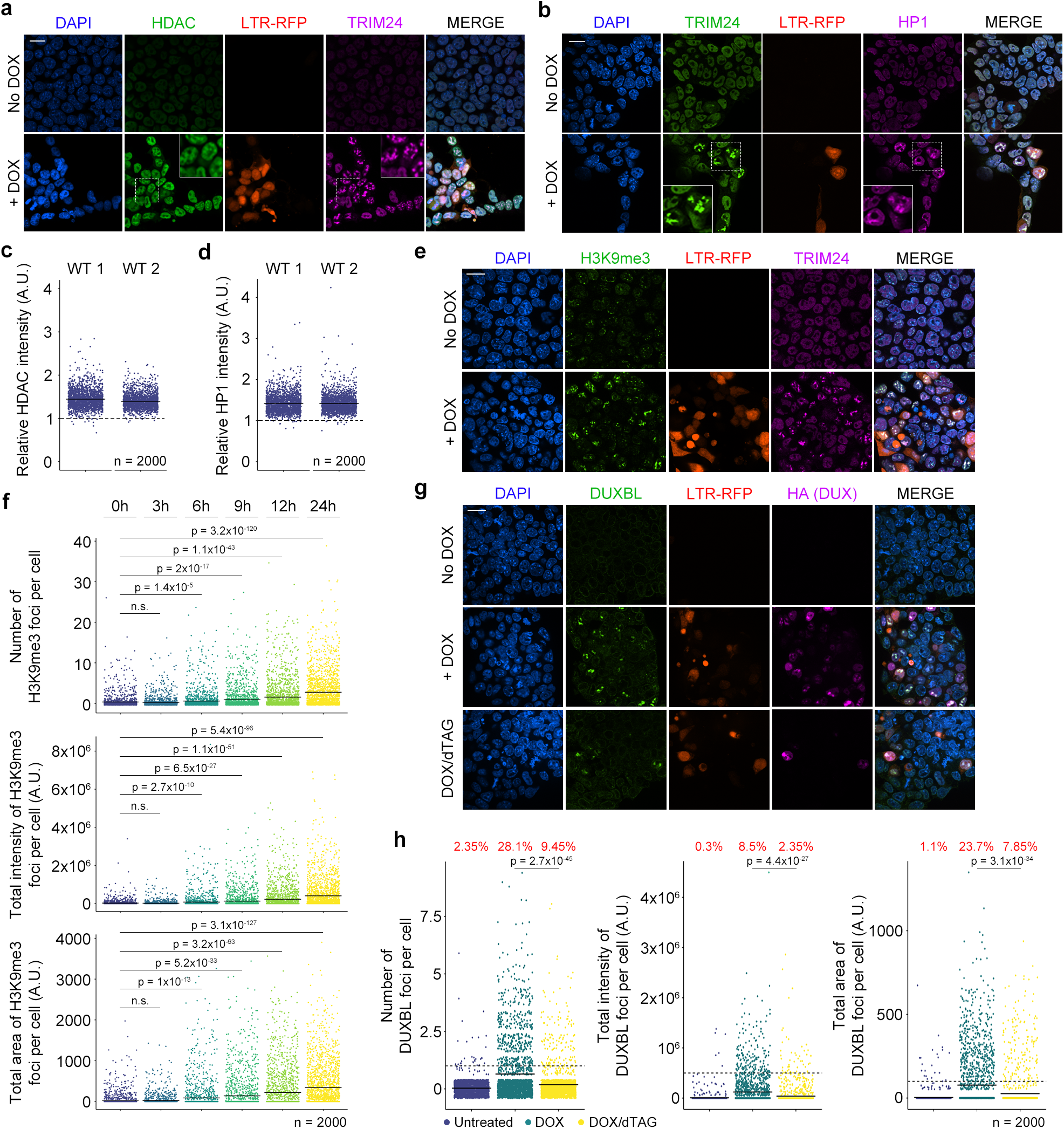
DUXBL and TRIM24/TRIM33 cooperatively silence the 2C-transcriptional program. **a, b)** Immunofluorescence analysis of HDAC (a) and HP1 (b) in untreated or DOX-treated LTR-RFP reporter ESC^DUX^. DAPI was used to visualize nuclei. Scale bars, 20 μm. Dashed regions are zoomed-in in the insets. Two independent experiments using two ESC^DUX^ clones were performed but one representative experiment is shown. **c, d)** Plot showing the ratio for HDAC (c) and HP1 (d) between the fluorescence intensity found at TRIM24 foci and the mean nuclear intensity per cell detected in DOX-treated LTR-RFP reporter ESC^DUX^. Center lines indicate mean values. n=2000. Two independent experiments using two ESC^DUX^ clones were performed but one representative experiment is shown. **e)** Immunofluorescence analysis of H3K9me3 in untreated or DOX-treated LTR-RFP reporter ESC^DUX^. DAPI was used to visualize nuclei. Scale bars, 20 μm. Three independent experiments using two ESC^DUX^ clones were performed but one representative experiment is shown. **f)** High-throughput imaging quantification of the number of H3K9me3 foci (upper panel), the total intensity of H3K9me3 foci (middle panel) and the total area of H3K9me3 foci (lower panel) per cell in untreated or DOX-treated LTR-RFP reporter ESC^DUX^. Center lines indicate mean values. n=2000; Relevant p values are shown from one-tailed unpaired *t*-tests. Three independent experiments using at least two ESC^DUX^ clones were performed but one representative experiment is shown. **g)** Immunofluorescence analysis of DUXBL in untreated or DOX-treated ESC^DUX-FKBP^. Treatments include DOX for 24 hours or DOX for 16 hours plus 8 hours with dTAG compounds. DAPI was used to visualize nuclei. Scale bar, 20 μm. Three independent experiments were performed but one representative experiment is shown. **h)** High-throughput imaging quantification of the number of DUXBL foci (left panel), the total intensity of DUXBL foci (middle panel) and the total area of DUXBL foci (right panel) per cell in ESC^DUX-FKBP^ treated as in (g). Center lines indicate mean values. n=2000; Relevant p values are shown from one-tailed unpaired *t*-tests. Percentages above the threshold (dotted line) are indicated. Three independent experiments were performed but one representative experiment is shown.

**Fig. 7:**
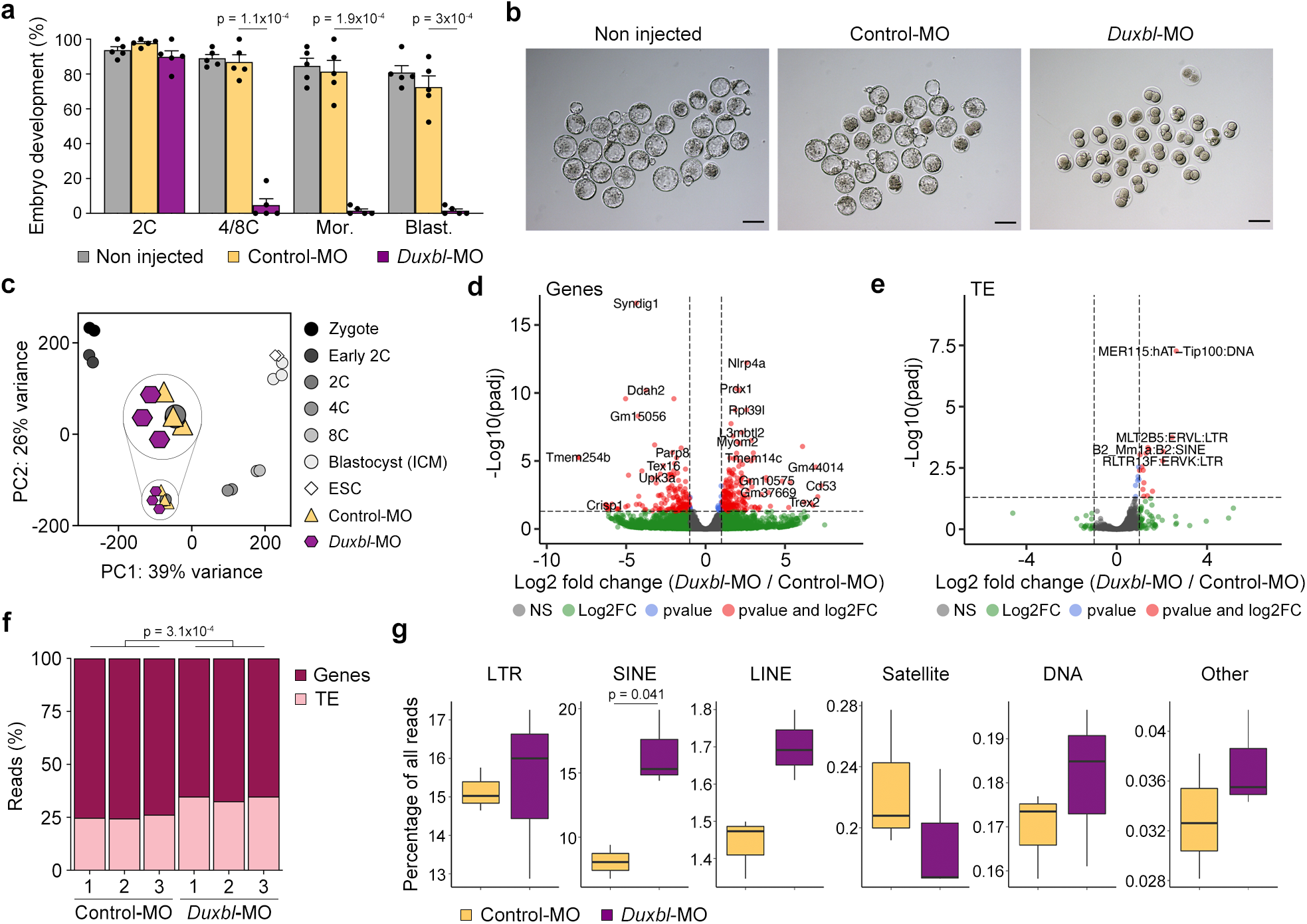
DUXBL is essential for development past the totipotent stage in mouse embryos. **a)** Plot summarizing five independent experiments with a total of 25-45 microinjected zygotes per group [non-microinjected, control morpholino (Control-MO), or *Duxbl* morpholino (*Duxbl* -MO)-injected zygotes] per experiment. The percentage of embryos reaching each embryo stage is shown. Mor: morula, Blast: blastocyst. p values are shown from one-tailed unpaired *t*-tests. **b)** Images from a representative microinjection experiment performed. Embryos were imaged three days after microinjection. Scale bar, 100 μm. **c)** PCA plot of RNA-seq data from control or *Duxbl* MO-injected zygotes collected at the late 2C-stage (three replicates per condition are shown) together with an available dataset of samples that evaluate temporal transcriptional dynamics during early mouse development^28^. The inset shows a zoomed-in section of the 2C samples. **d, e)** Volcano plots showing differentially expressed genes (d) and TE (e) between MO and control-injected zygotes collected at the late 2C-stage. Relevant genes and TE are highlighted. **f)** Plot showing the percentage of total reads mapping to genes or TE from samples described in (d, e). p value is shown from one-tailed unpaired *t*-test. **g)** Box and whisker plots showing the percentage of total reads mapping to different classes of TE from samples described in (d, e). p value is shown from one-tailed unpaired *t*-test. Center line indicates the median, box extends from the 25^th^ to 75^th^ percentiles and whiskers show Min to Max values.

## DISCUSSION

Upon exit from totipotency and transition to the 4C-embryo stage, the 2C-associated transcriptional program needs to be quickly and efficiently decommissioned to allow development to proceed. Our work suggests a model where DUX-mediated transcriptional activation promotes the expression of DUXBL and its recruitment along with the TRIM24/33 complex to DUX-bound regions to facilitate gene and MERVL silencing. We show that DUX induction of the *Duxbl* gene leads to two different DUXBL protein forms, DUXBL^SM^ and DUXBL^LG^, expressed at similar levels *in vivo*. Importantly, endogenous expression of DUXBL^SM^ is sufficient to compensate for the loss of DUXBL^LG^. Nevertheless, it is conceivable that DUXBL^SM^ and DUXBL^LG^ perform exclusive functions during the exit from totipotency. Indeed, we identified specific protein interactors for each DUXBL protein form, suggesting they might also exert unique functions. In support of our observations, activation of TE promoters can generate alternative protein isoforms during pre-implantation development^46^. The generation of different protein forms from the same gene, commonly affecting the N-terminal region, allows for the expansion of new protein functions needed during development. Further work will be necessary to dissect the precise role of each DUXBL isoform.

During ZGA, MERVL elements undergo extensive demethylation allowing LTRs to act as alternative promoters of 2C-associated protein-coding genes. Recent results have unveiled an epigenetic switch from DNA methylation to H3K9me3 deposition to regulate retrotransposon expression including MERVL elements^44,47^. Interestingly, DUX may contribute to the establishment of H3K9me3 regions as *Dux*-deficient embryos showed a significant reduction of H3K9me3 deposition^44^. Our data show that DUX-induced chromatin opening and MERVL activation leads to the formation of DUX-associated nuclear foci and the recruitment of the TRIM24/33/DUXBL complex. Indeed, ERV derepression in ESC leads to the formation of transcriptional condensates associated with transcribing ERV loci^48^. At these foci, TRIM24/33 further recruit HDAC proteins and HP1 promoting heterochromatin formation. We propose that in this genomic context DUXBL functions as a DUX-dependent dominant negative by competing with DUX for the same binding sites. Indeed, although DUXBL only has access to DUX-bound regions in the presence of DUX, overexpression of DUXBL is sufficient to overcome the proper activation of the 2C-transcriptional program. The increase of local DUXBL, further promoted by TRIM24-mediated recruitment, might displace DUX and associated HATs, ensuring transcriptional silencing and decreased chromatin accessibility. In summary, H3K9me3 establishment and silencing of the 2C-associated program seems facilitated by a negative feedback loop following DUX expression suggesting that a decreased activation of the 2C transcriptional program results in defective establishment of silencing signals.

We showed that DUXBL is required to exit from the totipotent 2C embryo stage. *Duxbl*-knockdown embryos undergo a 2C arrest characterized by extensive gene dysregulation and increased expression of TEs, particularly SINE B2 repeats. Importantly, upregulation of SINE B2 repeats is associated with detrimental embryonic development^49^. We did not detect specific upregulation of the 2C-transcriptional program in *Duxbl*-knockdown embryos likely due to incomplete DUXBL depletion or to the timing when samples were collected to perform RNA-seq. Indeed, the 2C-transcriptional program undergoes silencing during the transition to the 4C stage.

Finally, the *Duxbl* gene is only found in the mouse and rat genomes and raises the question of whether related genes present in other mammals could serve a similar function during embryonic development. Interestingly, DUXB, a gene found in all major mammalian lineages, has been proposed as a DUXBL paralog^24^ as they both share a similar gene structure, high homeodomain identity and are contained in a syntenic genomic region^24^. In mouse and rat, exons four and five of a DUXB-type gene have been identified, while in primates, decaying DUXBL pseudogenes were also found. This suggests an evolutionary pattern of reciprocal loss and retention of these two genes in different mammalian lineages with a likely similar function^24^. Nevertheless, we cannot exclude the possibility that other members of the DUX family, including DUXA or DUXC, could also perform some functions mediated by DUXBL in different species.

## METHODS

### One-cell embryo collection

Six to twelve-week-old NSA (CF-1) female mice (Envigo, Indianapolis, IN) were injected intraperitoneally with 7.5 IU eCG, followed 48 h later by 7.5 IU hCG and mating to B6SJLF1/J males (Jackson Laboratory, Bar Harbor, ME). Females were sacrificed, and 1-cell embryos were flushed from the oviducts of superovulated females 20 h post-hCG. The collection medium was MEM, HEPES supplemented with 0.1% PVA (MEM/PVA). Cumulus cells were removed by treatment with 1 mg/ml hyaluronidase in MEM/PVA. Embryos were transferred to KSOM at 37ºC in an atmosphere of 5% CO2, 5% O2 and 90% N2 until microinjection. All the animal work included here was performed in compliance with the NIH Animal Care & Use Committee (ACUC) Guideline for Breeding and Weaning.

### Oocyte collection

Germinal vesicle-intact (GV) oocytes were collected from 6-12-week-old female CF-1 mice 44-48 h after injection with 7.5 IU eCG. Females were sacrificed, ovaries were placed in MEM/PVA containing 2.5 µM milrinone, and antral follicles were obtained by poking with 27G needles. Cumulus-enclosed oocytes were collected, and cumulus cells were removed by pipetting up and down through a narrow bore capillary. Oocytes were placed in MEM*α* containing 5% fetal bovine serum (FBS) and 2.5 µM milrinone at 37ºC in a humidified atmosphere of 5% CO2 until microinjection. *In vitro* fertilization was performed as previously described^50^.

### Oocyte and 1-cell embryo microinjection

Oocyte and 1-cell embryo microinjections were performed using a Femtojet injector and TransferMan 4r micromanipulators (Eppendorf, Enfield, CT) on a Leica DMI 6000B inverted microscope. Oocytes/embryos were injected in the cytoplasm with 5-10 pl of morpholino oligonucleotides (MO), siRNAs, or a combination of siRNAs and MO. Morpholinos were injected at 1 mM concentration; siRNAs were injected at 50 µM concentration. The following MO were used: *Duxbl*-MO: 5’-ATCCCAAGAGGCCCAAGAGAGCTGT-3’, *Duxbl*-MO2: 5’-TAGAGCCCAGAGTGGCCCACCCATT-3’; standard control-MO: 5’-CCTCTTACCTCAGTTACAATTTATA-3’. The following siRNAs were used: *Duxbl* Silencer Select Pre-designed siRNA ID# s114932, Silencer Select Negative Control No. 1 (Thermo Fisher, Waltham, MA). Following microinjection, oocytes were put in the incubator in MEM*α* containing 5% FBS and 2.5 µM milrinone for 3-4 h and then in vitro matured for 16 h in MEM*α* + FBS. Embryos were placed back in the incubator and cultured for 4 days in KSOM in a humidified atmosphere of 5% CO2, 5%O_2_ and 90% N_2_.

### Cell culture

Wild-type (R1, G4, E14) ESC, ESC^DUX^ and ESC^CTCF-AID^ (ID: EN52.9.1) were grown on 0.1% gelatin-coated plates or alternatively on a feeder layer of growth-arrested MEFs in high-glucose DMEM (Gibco) supplemented with 15% FBS, 1:500 LIF (made in house), 0.1 mM non-essential amino acids, 1% glutamax, 1mM Sodium Pyruvate, 55 mM β-mercaptoethanol, and 1% penicillin/streptomycin (all from ThermoFisher Scientific) at 37°C and 5% CO2. Cells were routinely passaged with Trypsin 0.05% (Gibco). Media was changed every other day and passaged every 2-3 days. HEK293T (American Type Culture Collection) cells were grown in DMEM, 10% FBS, and 1% penicillin/streptomycin. Generation of infective lentiviral particles and ESC infections were performed as described^18^.

To generate DUXBL, TRIM24 and TRIM33 knockout ESC lines, we independently infected a suspension of ESC with lentiviral supernatants, generated by using LentiCRISPRv2 (gift from Feng Zhang, 52961, Addgene), encoding sgRNAs designed to target the corresponding protein (Supplementary Table 8 for sgRNA sequences).

To generate DOX-inducible DUXBL^LG^ and DUXBL^SM^ ESC lines as well as ESC^DUX-FKBP^ lines or ESC^DUX-2XHA^ lines, the corresponding sequences were PCR amplified from cDNA including a 3XFLAG for DUXBL^LG^ or an HA-tag for DUXBL^SM^ and subcloned by Gibson assembly into the plasmid PB-TRE-dCas9-VPR (Gift from George Church, 63800, Addgene), after removing the dCas9-VPR insert. DOX-inducible PiggyBac plasmids together with a plasmid encoding for a supertransposase were co-transfected in ESC using jetPRIME (PolyPlus transfection) and further selected with Hygromycin (200ug/ml) for one week. (Supplementary Table 8). To induce degradation of the DUX-FKBP, ESC^DUX-FKBP^ lines were treated with a combination of 500 nM dTAGv and dTAG13.

To generate DUXBL-BR1, BR2 and BR3 ESC^DUX^ reporter cell lines, the corresponding DUX-bound sequences were PCR amplified from mouse genomic DNA and subcloned in a PiggyBac plasmid upstream of an GFP coding region to generate the corresponding reporter (Supplementary Table 8). ESC^DUX^ carrying the LTR-RFP reporter were co-transfected with each of the BR-reporters together with a plasmid encoding for a supertransposase using jetPRIME (PolyPlus transfection) and further selected with Puromycin (1ug/ml) for one week.

To generate DUXBL^iRFP702^-knock-in ESC^DUX^ lines, we made a targeting construct using the backbone from the CTCF-mAC donor (Gift from Masato Kanemaki, 140646, Addgene) and containing the sequence Duxbl^Exon 5^-iRFP702-Duxbl^3’UTR^ after amplifying the corresponding sequences by PCR followed by Gibson assembly (New England Biolabs) (Supplementary Table 8). In addition, we also used a targeting construct to target *Oct4* with an IRES-GFP cassette (Gift from Rudolf Jaenisch, 21547, Addgene). DUXBL is not expressed in ESC and thus, to identified targeting events in *Duxbl*, ESC^DUX^ were co-transfected using jetPRIME (PolyPlus transfection) with both targeting vectors as well as with LentiCRISPRv2 encoding sgRNAs designed to target the sequences surrounding the stop codons of *Oct4* and *Duxbl*. Co-targeting events are common and therefore, GFP^+^ ESC^DUX^ were sorted on a BD FACSAria Fusion instrument and tested for iRFP702 expression upon DUX induction. Post-sort quality control was performed for each sample (Extended data Fig. 9).

### Cell sorting

For cell sorting experiments, trypsinized live OCT4-IRES-GFP ESC with the specific gene targeting (DUXBL) were sorted based on GFP fluorescence levels on a BD FACSAria Violet instrument (Extended data Fig. 9). Post-sort quality control was performed for each sample (Extended data Fig. 9).

### Immunofluorescence

Cells were fixed in 4 % Paraformaldehyde (PFA, Electron Microscopy Sciences) for 10 min at RT followed by 10 min of permeabilization using the following permeabilization buffer (100 mM Tris-HCl pH 7.4, 50 mM EDTA pH 8.0, 0.5 % Triton X-100). The following primary antibodies were incubated overnight: TRIM24 (1:200, TA802797, Origene), TRIM33 (1:200, A301-060A, Bethyl Laboratories), DUXBL (1:500, custom antibody, GenScript), HDAC1 (1:100, D5C6U, #34589, Cell Signaling), HP1 (1:200, #2616, Cell Signaling), HA (1:500, C29F4, #3724, Cell Signaling), HA (1:100, 6E2, #2367, Cell Signaling), H3K9me3 (1:500, D4W1U, #13969, Cell Signaling), FLAG (1:500, F1804, Sigma Aldrich). Corresponding Alexa Fluor 488 Chicken anti-Rabbit IgG (H+L) (Thermo Fisher Scientific, Cat# 31431), Alexa Fluor 488 Goat anti-Mouse IgG (H+L) (Thermo Fisher Scientific, Cat# A-11001), Alexa Fluor 647 Chicken anti-Rabbit IgG (H+L) (Thermo Fisher Scientific, Cat# A-21443) or Alexa Fluor 647 Chicken anti-Mouse IgG (H+L) (Thermo Fisher Scientific, Cat# A-21463) secondary antibodies were used to reveal primary antibody binding (1:1000). DNA was stained using DAPI (4′,6-diamidino-2-phenylindole). Images were acquired using a Nikon SoRa spinning disk microscope (CSU-W1). Image processing was done using Nikon’s NIS-Element.

### High throughput imaging (HTI)

A total of 10,000-20,000 ESC (depending on the experiment and on the specific ESC line) were plated on gelatinized μCLEAR bottom 96-well plates (Greiner Bio-One, 655087). ESC were treated with DOX (different concentrations in the range from 10–1000 ng/ml depending on the experiment and on the specific ESC line), 2.5 µM PlaB, 0.53 µM RA or 500 mM IAA as indicated before fixation with 4% PFA in PBS for 10 minutes at room temperature.

Images were automatically acquired either using a CellVoyager CV7000 or a CellVoyager CV8000 high throughput spinning disk confocal microscope (Yokogawa, Japan). Each condition was always performed in triplicate wells and at least 9 different fields of view (FOV) were acquired per well. Image analysis was performed using the Columbus Image Data Storage and Analysis system (PerkinElmer). Nuclei were segmented based on DAPI staining. Detection of nuclear foci was performed on their respective fluorescence channels using the Find Spots module. Various measurements were calculated over the nuclear masks, including mean fluorescence intensities and (when applicable) the number, intensity, area, and colocalization of foci. Cell-level data was exported from Columbus as text files, then analyzed and plotted using R version 4.1.0. When analyzing HTI data, we considered statistically significant those samples that when compared showed an unpaired one or two-tail t-test with a p-value of at least 0.05 or lower.

### Live cell imaging

A total of 40,000 ESC^DUX^ stably transfected with LTR-RFP and BR1, BR2 or BR3-eGFP PiggyBAC constructs were plated in gelatin-coated m-Slide 8 wells plates (80826, Ibidi) and imaged untreated or DOX-treated every 20 minutes for a total time of 15 hours. To examine 2C-like conversion dynamics, 40,000 WT or DUXBL^KO^ ESC, stably transfected with LTR-RFP construct, were imaged every 30 min for a total time of 48hrs. To visualize DUXBL foci formation, a total of 40,000 000 ESC^DUX^ stably transfected with LTR-RFP and endogenously tagged with a miRFP702 fluorescent protein at the *Duxbl* loci constructs were imaged in untreated or DOX-treated ESC^DUX^ every 1 hour and 30 minutes for a total time of 14 hours. All images were acquired using the Nikon SoRa spinning disk confocal microscope equipped with 20x plan-apochromat objective lenses (N.A. 0.75 and 0.8, respectively) and stage top incubators to maintain temperature, humidity and CO2 (Tokai Hit STX and Okolab Bold Line, respectively). Image processing was done using Nikon’s NIS-Element. In case of the experiments analyzing DUXBL foci or BR-GFP reporter ESC lines, images were denoised using Nikon’s NIS-Element denoise AI.

### Western blot

Cells were trypsinized and lysed in 50 mM Tris pH 8, 8 M Urea (Sigma) and 1% Chaps (Millipore) followed by 30 min of shaking at 4°C. A total of 20 μg of extracts were run on 4%-12% NuPage Bis-Tris Gel (Invitrogen) and transferred onto Nitrocellulose Blotting Membrane (GE Healthcare). Transferred membranes were incubated with the following primary antibodies overnight at 4°C: DUXBL (1:1000, Custom antibody, GenScript), ZSCAN4C (1:500, AB4340, Millipore Sigma), TRIM24 (1:1000, TA802797, Origene), TRIM33 (1:1000, A301-060A, Bethyl Laboratories), HA (1:1000, C29F4, #3724, Cell Signaling), HA (1:1000, 6E2, #2367, Cell Signaling), H2A (1:1000, ab18255, Abcam), FLAG (1:1000, F1804, Sigma Aldrich), Tubulin (1:50000, T9026, Sigma-Aldrich). The next day the membranes were incubated with HRP-conjugated secondary antibodies Goat anti-Rabbit IgG (H+L) (1:5000; Thermo Fisher Scientific, Cat# 31466) or Goat anti-Mouse IgG (H+L) (1:5000; Thermo Fisher Scientific, Cat# 31431) for 1 hour at room temperature. Membranes were developed using SuperSignal West Pico PLUS or SuperSignal West Femto Maximum sensitivity (Thermo Scientific).

### RNA extraction, qPCR and RACE analysis

Isolation of total RNA and cDNA synthesis was performed by using the Isolate II RNA Mini Kit (Bioline) and SensiFAST cDNA Synthesis Kit (Bioline), respectively. Quantitative real time PCR was performed with PowerUp SYBR Master mix in a QuantStudio 6 Pro system. Expression levels were normalized to GAPDH. For a complete primer list see Supplementary Table 8. When analyzing quantitative real time PCR data, we considered statistically significant those samples that when compared showed an averaged of two-fold difference in overall gene expression and an unpaired two-tail t-test with a p-value of at least 0.05 or lower. For RACE analysis we used the SMARTer® RACE 5’/3’ Kit (Takara) according to manufacturer’s recommendations (see Supplementary Table 8 for the list of primers used).

### Immunoprecipitation

To obtain nuclear extracts, ESC were trypsinized, washed with ice-cold phosphate-buffered saline (PBS), resuspended in 1 volume of ice-cold hypotonic lysis buffer (10 mM HEPES pH 7.9, 10 mM KCl, 0.1 mM EDTA containing protease inhibitors) and incubated on ice for 10 minutes. Next, 1/10 volume of 1% IGEPAL CA630 (Sigma, Merck) was added to the samples and incubated for additional 3 minutes at room temperature. After that, cells were briefly vortexed, and the cytosolic fraction was obtained by centrifugation for 5 min at 2,500 *g*. The nuclear pellet was resuspended in 1 volume of high-salt-concentration extraction buffer (20 mM HEPES pH 7.9, 0.6 M NaCl, 1 mM EDTA containing protease inhibitors) followed by incubation with shaking at 4°C for 10 min, sonicated briefly (until clear) and incubated with shaking at 4°C for at least 30 min more. The nuclear extract was obtained by collecting the supernatant after centrifugation for 5 min at 16,000 *g*. Protein concentration was determined using the Bradford assay.

A total of 1 or 2 mg of nuclear extract was diluted in binding buffer (25 mM Tris pH 7.9, 200 mM NaCl and 0.5 mM EDTA) and centrifuged for 5 min at 16,000 g at 4°C. Protein G Dynabeads (Invitrogen, Thermo Fisher Scientific) were washed twice with binding buffer and then incubated with 3 µg per milligram of protein of DUXBL antibody, FLAG antibody, HA antibody or a non-specific IgG in the presence of 0.5 µg/ml BSA. Alternatively, anti-FLAG-agarose beads (DYKDDDDK Fab-TrapTM Agarose, FFA, Proteintech) were also used to pull down FLAG-tagged proteins. The antibody-bound beads were washed 5 times with binding buffer and incubated with the cleared supernatant overnight (for Protein G Dynabeads) or for 1 h (for anti-FLAG-agarose beads) at 4°C. The beads were then washed five times with binding buffer with 0.05% IGEPAL CA630 (Sigma, Merck). One tenth of the beads were eluted in loading buffer for Western blot purposes and the rest was processed for Mass Spectrometry analysis.

### Mass spectrometry analysis

Samples were solution digested with trypsin using S traps (Protifi), following the manufacturer’s instructions. Briefly, proteins were denatured in 5% SDS, 50 mM triethylammonium bicarbonate (TEAB) pH 8.5. They were next reduced with 5 mM Tris(2-carboxyethyl)phosphine (TCEP) and alkylated with 20 mM iodoacetamide. The proteins were acidified to a final concentration of 2.5% phosphoric acid and diluted into 100 mM TEAB pH 7.55 in 90% methanol and loaded onto the S-traps, washed four times with 100 mM TEAB pH 7.55 in 90% methanol, and digested with trypsin overnight at 37 °C. Peptides were eluted from the S-trap using 50 mM TEAB pH 8.5; 0.2% formic acid in water; and 50% acetonitrile in water. These elutions were pooled and dried by lyophilization.

Dried peptides were resuspended in 5% acetonitrile, 0.05% TFA in water for mass spectrometry analysis on either an Obitrap Fusion Tribrid (Thermo Scientific) or an Orbitrap Exploris 480 (Thermo Scientific) mass spectrometer. The peptides were separated on a 75 µm x 15 cm, 3 µm Acclaim PepMap reverse phase column (Thermo Scientific) at 300 nL/min using an UltiMate 3000 RSLCnano HPLC (Thermo Scientific) and eluted directly into the mass spectrometer. For analysis in the Fusion, parent full-scan mass spectra collected in the Orbitrap mass analyzer set to acquire data at 120,000 FWHM resolution and HCD fragment ions detected in the ion trap. For analysis in the Exploris 480, parent full-scan mass spectra acquired at 120,000 FWHM resolution and product ion spectra at 15,000 resolution.

Proteome Discoverer 2.4 (Thermo) was used to search the data against the murine database from Uniprot using SequestHT. The search was limited to tryptic peptides, with maximally two missed cleavages allowed. Cysteine carbamidomethylation was set as a fixed modification, with methionine oxidation as a variable modification. The precursor mass tolerance was 10 ppm, and the fragment mass tolerance was 0.6 Da for data obtained on the Fusion and 0.02 Da for data obtained on the Exploris 480. The Percolator node was used to score and rank peptide matches using a 1% false discovery rate. Label-free quantitation of extracted ion chromatograms from MS1 spectra was performed using the Minora node in Proteome Discoverer.

### CUT&RUN protocol

The CUT&RUN protocol was performed as described^18^. In brief, 500,000 trypsinized ESC were washed a total of three times with Wash Buffer (20 mM HEPES-KOH pH 7.5, 150 mM NaCl, 0.5 mM spermidine, Roche complete Protease Inhibitor tablet EDTA free) and bound to activated Concanavalin A beads (Polysciences) for 10 minutes at room temperature. Cell permeabilization was done in Digitonin Buffer (0.05 % Digitonin and 0.1% BSA in Wash Buffer) followed by incubation with 4µl of the antibody against DUXBL (TA331435, Origene), 1µl of the antibody against FLAG (F1804, Sigma Aldrich) or 2µl of the antibody against H3K27ac (ab4729, Abcam) at 4°C for 2 hours. For negative controls, we used a Guinea Pig anti-Rabbit IgG (ABIN101961, Antibodies-online). After antibody incubation, cells were washed with Digitonin Buffer, followed by incubation with purified hybrid protein A-protein G-Micrococcal nuclease (pAG-MNase) at 4°C for 1 hour. Samples were washed again in Digitonin Buffer, resuspended in 150 μl Digitonin Buffer and equilibrated to 0°C on ice water for 5 minutes. To start with MNase cleavage, 3 μl of 100 mM CaCl_2_ was added to cells and after 1 hour of digestion, reactions were stopped with the addition of 150 μl 2x Stop Buffer (340 mM NaCl, 20 mM EDTA, 4 mM EGTA, 0.02 % Digitonin, 50 μg/ml RNase A, 50 μg/ml Glycogen). Samples were incubated at 37°C for 10 minutes to release DNA fragments and centrifuged at 16,000 g for 5 minutes. Supernatants were collected and a mix of 1.5 μl 20% SDS / 2.25 μl 20 mg/ml Proteinase K was added to each sample followed by incubation at 65°C for 35 minutes. DNA was precipitated with ethanol and sodium acetate and pelleted by centrifugation at 4°C, washed, air-dried and resuspended in 10 μl 0.1x TE.

### Library preparation and sequencing

For all experiments, we used half of the precipitated DNA obtained from CUT&RUN to prepare Illumina compatible sequencing libraries as described^18^. In brief, end-repair was performed in 50 μl of T4 ligase reaction buffer, 0.4 mM dNTPs, 3 U of T4 DNA polymerase (NEB), 9 U of T4 Polynucleotide Kinase (NEB) and 1 U of Klenow fragment (NEB) at 20°C for 30 minutes. Following this step, end-repair reaction was cleaned using AMPure XP beads (Beckman Coulter) and eluted in 16.5 μl of Elution Buffer (10 mM Tris-HCl pH 8.5) followed by A-tailing reaction in 20 μl of dA-Tailing reaction buffer (NEB) with 2.5 U of Klenow fragment exo-(NEB) at 37°C for 30 minutes. The total volume of the A-tailing reaction was mixed with Quick Ligase buffer 2X (NEB), 3000 U of Quick Ligase (NEB) and 10 nM of annealed adaptor (Illumina truncated adaptor) in a final volume of 50 μl and incubated for 20 min at room temperature. The adaptor was prepared by annealing the following HPLC-purified oligos: 5′-Phos/GATCGGAAGAGCACACGTCT-3′ and 5′-ACACTCTTTCCCTACACGACGCTCTTCCGATC∗T-3′ (∗phosphorothioate bond). Ligation was stopped by adding 50 mM of EDTA, cleaned with AMPure XP beads and eluted in 14 μl of Elution Buffer. All volume was used for PCR amplification in a 50 μl reaction with 1 μM primers TruSeq barcoded primer p7, 5′-CAAGCAGAAGACGGCATACGAGATXXXXXXXXGTGACTGGAGTTCAGACGTGTGCTCTTCC GATC∗T-3′ and TruSeq barcoded primer p5 5′-AATGATACGGCGACCACCGAGATCTACACXXXXXXXXACACTCTTTCCCTACACGACGCTC TTCCGATC*T-3′ (∗ represents a phosphothiorate bond and XXXXXXXX a barcode index sequence), and 2X Kapa HiFi HotStart Ready mix (Kapa Biosciences). PCR amplification was performed with the following program: 45 s at 98°C followed by 15 cycles of 15 s at 98°C, 30 s at 63°C, 30 s at 72°C and a final 5 min extension at 72°C. PCR reactions were cleaned with AMPure XP beads (Beckman Coulter), run on a 2% agarose gel and a band of 300bp approximately was cut and gel purified using QIAquick Gel Extraction Kit (QIAGEN). Library concentration was determined with KAPA Library Quantification Kit for Illumina Platforms (Kapa Biosystems). Sequencing was performed on the Illumina NextSeq550 (75bp pair-end reads) or on the Illumina NextSeq2000 (50bp pair-end reads).

### CUT&RUN data processing

Data were processed using a modified version of CUT&RUNTools as previously reported^18^. Reads were adapter trimmed using fastp v.0.20.0. An additional trimming step was performed to remove up to 6bp adapter from each read. Next, reads were aligned to the mm10 genome using bowtie2 with the ‘dovetail’ and ‘sensitive’ settings enabled. Peaks were called using macs2^51^ with a q-value cutoff of <0.01. For broad peaks calling, SICER v1.1 was used with parameters “redundancy threshold”=100, “window-size”=300, “gap size”=600 and E-value= 100. Signal tracks normalized to 1X coverage (reads per genomic coverage, RPGC) were generated using the ‘bamCoverage’ utility from deepTools^52^ with parameters bin-size=25, smooth length=75, and ‘center_reads’ and ‘extend_reads’ options enabled. For motif analysis of DUXBL peaks, FIMO^53^ from the MEME suite was used to scan either the entire peak sequence or the 300bp of sequence surrounding peak summits using the predicted DUXBL motif from the JASPAR database (https://jaspar.genereg.net/matrix/UN0671.1/).

### RNA-seq libraries

RNA-seq libraries were prepared using NEBNext Ultra II Directional RNA Library Prep Kit for Illumina (New England Biolabs, NEB) and NEBNext rRNA Depletion Kit (Human/Mouse/Rat) (NEB) according to the manufacturer’s protocol. Sequencing was performed on the Illumina NextSeq500 (75bp pair-end reads).

### RNAseq data processing and batch correction

Fastq files for publicly available RNAseq experiments were downloaded from SRA. RNAseq reads were adapter trimmed using cutadapt v1.18^54^. Trimmed reads were aligned to either the mm10 genome with the GENCODE transcriptome annotation version M25 or the mm39 genome with the GENCODE transcriptome annotation M28 using STAR v2.7.6a^55^. Transcript and transposable element expression was quantified using TEcount from tetoolkit v2.1.4^56^. Differential expression analysis was performed using the DESeq2 package^57^. To compare the RNAseq experiments with public data, batch correction was performed. Gene counts across samples were first quantile-normalized using the limma package. Batch correction was then performed on quantile-normalized counts using COMBAT from the sva package.

### DUXBL custom antibody

To generate a custom rabbit polyclonal antiserum (GenScript), a poly-histidine tagged DUXBL fragment (aminoacids 193-350) was expressed in E. coli and purified by GenScript protein department. Two New Zealand rabbits (rabbit No. R04775 and R04776) were immunized with three injections (200mg/animal) every other two weeks by conventional protocol. Freund’s adjuvant was added to the immunogen. Seven days following the 3rd immunization, the titer of antiserums was tested by ELISA. Based on in-house testing results, the rabbits were sacrificed according to animal welfare principles and the final antiserum was purified by using antigen affinity column. Total IgG from pre-immune serum was used as negative control. 0.02% Sodium Azide was added in the final antiserums as preservative.

## ACKNOWLEDGMENTS

We thank principal investigators from the Laboratory of Genome Integrity for helpful comments and discussion on this work. We also thank Mariam Malik, David Goldstein and the CCR Genomics Core for sequencing support, Ferenc Livak and the CCR Flow cytometry Core, Gianluca Pegoraro and the High-Throughput Imaging Facility and Michael Kruhlak and the Microscopy Core for experimental support. Research in S.R. and C.J.W. laboratory is supported by the Intramural Research Program of the NIH. T.O. is supported by a postdoctoral fellowship of the Helen Hay Whitney Foundation.

## AUTHOR CONTRIBUTIONS

M.V-S., T.O. and S.R. conceived the study. T.O., M.V-S. designed, performed, and analyzed *in vitro* experiments. P.S. and V.S. performed microinjection and *in vivo* experiments. C.N.D. provided technical support. D.T. analyzed sequencing data. G.I.C. provided support with high-throughput microscopy imaging. B.S. analyzed live confocal microscopy data. T.K.M. and L.M.J. performed and analyzed Mass Spectrometry experiments. S.R. and C.J.W. supervised the study and wrote the manuscript with comments and help from all authors.

## DATA AVAILABILITY

The sequencing data generated in this study have been deposited in the Gene Expression Omnibus database under accession code GSE210892. Datasets obtained from publicly available sources include GSE98149, GSE95517, GSE45719, GSE85627, GSE66582 and GSE97304 (https://www.ncbi.nlm.nih.gov/geo/query/acc.cgi?acc=GSE98149/GSE95517/GSE45719/GSE85627/GSE66582andGSE97304, respectively). Additional data and/or reagents that support the findings of this study are available from the corresponding author upon reasonable request. Source data are provided with this paper.

