## Supplementary Material for "The homeobox transcription factor DUXBL controls exit from totipotency"

**EXTENDED DATA LEGENDS:**

**Extended Data Fig. 1: a)** High-throughput imaging quantification of GFP mean nuclear intensity in untreated or DOX-treated for 16 hours BR1-GFP reporter ESC$^{DUX}$. Center lines indicate mean values. n=537. **b)** Graph plots showing quantified mean intensity levels of RFP and GFP obtained from untreated and DOX-treated LTR-RFP reporter BR2- or BR3-GFP ESC$^{DUX}$. **c)** Schematic representation of the *Duxbl* RNA isoforms detected by RACE analysis in DOX-treated ESC$^{DUX}$. **d)** DNA sequences surrounding exon 2 of *Duxbl* showing the deletions detected in representative DUXBL$^{LG\text{-}KO}$ and DUXBL$^{KO}$ ESC$^{DUX}$ clones analyzed. Exon sequence from exon 2 is highlighted in bold. **e)** Schematic representation of the DUXBL$^{LG}$ and DUXBL$^{SM}$ protein products.

**Extended Data Fig. 2: a)** Time lapse microscopy experiment following DOX induction performed in LTR-RFP reporter WT and DUXBL$^{KO}$ ESC$^{DUX}$. Time since the addition of DOX is indicated. Scale bar, 10 µm. **b)** High-throughput imaging (HTI) quantification of RFP$^{+}$ cells in LTR-RFP reporter WT ESC and DUXBL$^{KO}$ ESC. Center lines indicate mean values. Percentages of RFP$^{+}$ cells above the threshold (dotted line) are indicated. n=2000; p value is shown from one-tailed unpaired *t*-test. Three independent experiments were performed but only one representative is shown. **c)** Unidimensional PCA plot of RNAseq data from untreated (two replicates each) or DOX-treated WT, DUXBL$^{LG\text{-}KO}$ and DUXBL$^{KO}$ ESC$^{DUX}$ together with WT DUX-expressing ESC from[5]. DUX-expressing ESC were sorted into GFP+ or GFP- based on the activation of the LTR-GFP reporter. **d)** Box and whisker plot showing normalized fold change (log2) expression of the 244 genes downregulated in DOX-treated DUXBL$^{KO}$ ESC$^{DUX}$ compared to DOX-treated WT ESC$^{DUX}$ during preimplantation development including ESC. RNAseq data obtained from[36]. Center line indicates the median, box extends from the 25$^{th}$ to 75$^{th}$ percentiles and whisker extends from the hinge to the largest or smallest value no further than 1.5-fold from the inter-quartile range. Data beyond the whiskers are considered outliers and plotted individually.

**Extended Data Fig. 3: a)** Western blot analysis of DUXBL (FLAG) performed in lysates from untreated or indole-3-acetic acid (IAA)/DOX-treated WT or ESC$^{CTCF\text{-}AID}$ expressing DUXBL$^{LG}$ (FLAG). Tubulin levels are shown as a loading control. **b)** Western blot analysis of CTCF performed in lysates from untreated or IAA-treated ESC$^{CTCF\text{-}AID}$. Tubulin levels are shown as a loading control. Note the increase in DUXBL expression on CTCF-depleted ESC. **c)** High-throughput imaging (HTI) quantification of $RFP^+$ cells in untreated or IAA/DOX-treated for 48 hours LTR-RFP reporter $ESC^{CTCF\text{-}AID}$ expressing $DUXBL^{LG}$. Center lines indicate mean values. Percentages of $RFP^+$ cells above the threshold (dotted line) are indicated. n=2000. Relevant p values are shown from one-tailed unpaired *t*-tests. **d)** High-throughput imaging quantification of $RFP^+$ cells in untreated or IAA/DOX-treated for 24 and 48 hours LTR-RFP reporter WT or $DUXBL^{KO}$ $ESC^{CTCF\text{-}AID}$. Center lines indicate mean values. Percentages of $RFP^+$ cells above the threshold (dotted line) are indicated. n=2000. Relevant p values are shown from one-tailed unpaired *t*-tests.

**Extended Data Fig. 4: a)** Predicted binding sites for DUX and DUXBL obtained from the Jaspar database. **b)** Venn diagrams showing the number of $DUXBL^{LG}$ peaks overlapping between those identified from CUT&RUN experiments performed with a DUXBL antibody (blue) and a FLAG antibody (yellow) in $ESC^{DUXBL\text{-}LG}$. **c)** Genome browser tracks corresponding to samples from (b) showing $DUXBL^{LG}$ occupancy at the indicated genomic regions revealing equivalent genomic distribution with both DUXBL and FLAG antibody. Input (IgG) is shown as reference control. **d)** CUT&RUN read density plot (RPGC) showing $DUXBL^{LG}$ and DUX enrichment at MERVL-int elements in DOX-treated $ESC^{DUXBL\text{-}LG}$. Data from DUX-expressing ESC was obtained from[5]. Input (IgG) is shown as reference control. **e)** Venn diagrams showing the number of $DUXBL^{LG}$ (left panel, blue) and DUX peaks (right panel, blue) overlapping with accessible regions as defined by ATAC-seq in ESC (yellow). ATAC-seq data was obtained from[35]. **f)** Genome browser tracks showing $DUXBL^{LG}$ and DUX occupancy as well as ATACseq signal at the indicated genomic regions in DOX-treated WT ESC expressing DUX or $DUXBL^{LG}$. Data from DUX-expressing ESC and ATACseq experiments were obtained from[5,35]. Input (IgG) is shown as reference control. **g)** Immunofluorescence analysis of $DUXBL^{LG}$ (FLAG) in untreated or DOX-treated LTR-RFP reporter $ESC^{DUXBL\text{-}LG/DUX}$. DAPI was used to visualize nuclei. Scale bar, 20 µm. **h)** Immunofluorescence analysis of $DUXBL^{LG}$ in untreated or DOX-treated $ESC^{DUXBL\text{-}LG}$. DAPI was used to visualize nuclei. Scale bar, 100 µm.

**Extended Data Fig. 5: a-c)** Western blot analysis of the indicated proteins performed with total protein extracts (input) and DUXBL immunoprecipitates (10% of total IP) obtained from untreated or DOX-treated $ESC^{DUXBL\text{-}LG}$ (a), untreated or DOX-treated $ESC^{DUX}$ (b) or untreated or DOX-treated $DUXBL^{LG\text{-}KO}$ $ESC^{DUX}$ (c). **d)** Volcano plot showing the enrichment of proteins obtained from endogenous DUXBL immunoprecipitation followed by mass spectrometry (IP-MS) analysis in two independent WT untreated or DOX-treated $ESC^{DUX}$. **e)** Volcano plot showing the enrichment of proteins obtained from endogenous DUXBL immunoprecipitation followed by mass spectrometry (IP-MS) analysis in two independent untreated or DOX-treated $DUXBL^{LG\text{-}KO}$ $ESC^{DUX}$ (these ESC only express $DUXBL^{SM}$ upon DUX expression).

**Extended Data Fig. 6: a)** Immunofluorescence analysis of DUX (HA) and endogenous TRIM24 in untreated or DOX-treated $ESC^{DUX\text{-}2XHA}$. DAPI was used to visualize nuclei. Scale bars, 20 µm. Two independent experiments were performed but one representative experiment is shown. **b)** Immunofluorescence analysis of endogenous TRIM24 in endogenous 2CLC observed in LTR-RFP reporter WT ESC. DAPI was used to visualize nuclei. Scale bars, 20 µm. Two independent experiments were performed but one representative experiment is shown. **c)** High-throughput imaging quantification of the number of TRIM24 foci (upper panel), the total intensity of TRIM24 foci (middle panel) and the total area of TRIM24 foci (lower panel) per cell in untreated or DOX-treated LTR-RFP reporter WT or $DUXBL^{KO}$ $ESC^{DUX}$. Center lines indicate mean values. n=2000; Relevant p values are shown from one-tailed unpaired *t*-tests. Percentages of cells above the threshold (dotted line) are indicated. Two independent experiments using at least two WT or $DUXBL^{KO}$ $ESC^{DUX}$ clones were performed but one representative experiment is shown. **d)** Western blot analysis of TRIM24 performed in WT and *Trim24*-knockout $ESC^{DUX}$. Tubulin levels are shown as a loading control. **e)** Western blot analysis of TRIM33 performed in WT and $TRIM33^{KO}$ $ESC^{DUX}$ lysates. Tubulin levels are shown as a loading control. **f)** High-throughput imaging quantification of the number of DUXBL foci (upper panel), the total intensity of DUXBL foci (middle panel) and the total area of DUXBL foci (lower panel) per cell in untreated or DOX-treated LTR-RFP reporter WT or $TRIM33^{KO}$ $ESC^{DUX}$. Percentages of cells above the threshold (dotted line) are indicated. Center lines indicate mean values. n=2000; Relevant p values are shown from one-tailed unpaired *t*-tests. Two independent experiments using at least two WT or $TRIM33^{KO}$ $ESC^{DUX}$ clones were performed but one representative experiment is shown. **g)** Western blot analysis of the indicated proteins performed with DUXBL immunoprecipitates obtained from untreated or DOX-treated WT and $TRIM24^{KO}$ $ESC^{DUX}$. H2A levels are shown as a loading control.

**Extended Data Fig. 7: a, b)** ChIPseq read density plot (RPGC) showing H3K9me3 enrichment at DUX-bound sites (a) and MERVL (b) elements occupied by $DUXBL^{LG}$ after DUX expression during embryonic development. Input (IgG) is shown as reference control. ChIP-seq data obtained from[44]. **c)** Graphs showing TRIM24 and H3K9me3 fluorescence intensity at TRIM24-foci at different times after the addition of DOX in ESC$^{DUX}$. Two independent experiments were performed but one representative experiment is shown. **d)** High-throughput imaging quantification of the number of DUX (HA) foci (upper panel), the total intensity of DUX (HA) foci (middle panel) and the total area of DUX (HA) foci (lower panel) per cell in untreated or DOX-treated ESC$^{DUX\text{-}FKBP}$. Treatments include DOX for 24 hours or DOX for 16 hours plus 8 hours with dTAG compounds. Percentages of cells above the threshold (dotted line) are indicated. Center lines indicate mean values; n=2000. Relevant p values are shown from two-tailed unpaired *t*-tests. Two independent experiments were performed but one representative experiment is shown. **e)** High-throughput imaging quantification of the number of TRIM24 foci (upper panel), the total intensity of TRIM24 foci (middle panel) and the total area of TRIM24 foci (lower panel) per cell in ESC$^{DUX\text{-}FKBP}$ treated as in (d). Center lines indicate mean values; n=2000. Relevant p values are shown from two-tailed unpaired *t*-tests. Percentages of cells above the threshold (dotted line) are indicated. Two independent experiments were performed but one representative experiment is shown.

**Extended Data Fig. 8: a)** Plot summarizing four independent experiments with a total of 25-45 microinjected zygotes per group (non-microinjected, control morpholino (MO), or a second morpholino (MO2)-injected zygotes) per experiment. The percentage of embryos reaching each embryo stage is shown. Mor: morula, Blast: blastocyst. p values are shown from one-tailed unpaired *t*-tests. **b)** Plot summarizing four independent experiments with a total of 25-45 microinjected zygotes per group (non-microinjected, control or *Duxbl* siRNAs-injected GV oocytes) per experiment. The percentage of embryos reaching each embryo stage is shown. Mor: morula, Blast: blastocyst**.** p values are shown from one-tailed unpaired *t*-tests. **c)** Relative fold change (log2) expression of *Duxbl* in uninduced or DOX-induced in ESC$^{DUX}$ transfected 36 hours prior to induction with the corresponding siRNAs. Reactions were performed by duplicate in two different ESC$^{DUX}$ lines. A representative experiment is shown. p values are shown from one-tailed paired *t*-tests. p value is shown from one-tailed unpaired *t*-test. **d)** Plot showing differential expression in different classes of TE from control or MO-injected zygotes collected at the late 2C-stage. **e)** Plot showing the percentage of SINE B2 reads compared to total TE reads from samples described in (d).

**Extended Data Fig. 9:** Flow cytometry plots generated from ESC$^{DUX}$ after performing the OCT4-IRES-GFP and DUXBL$^{iRFP702}$-knock-in targeting as described in Methods. Gating shows the actual GFP+ cells sorted for the experiment. Two independent ESC$^{DUX}$ lines were targeted and subclones isolated from them with similar results, but one representative is shown.

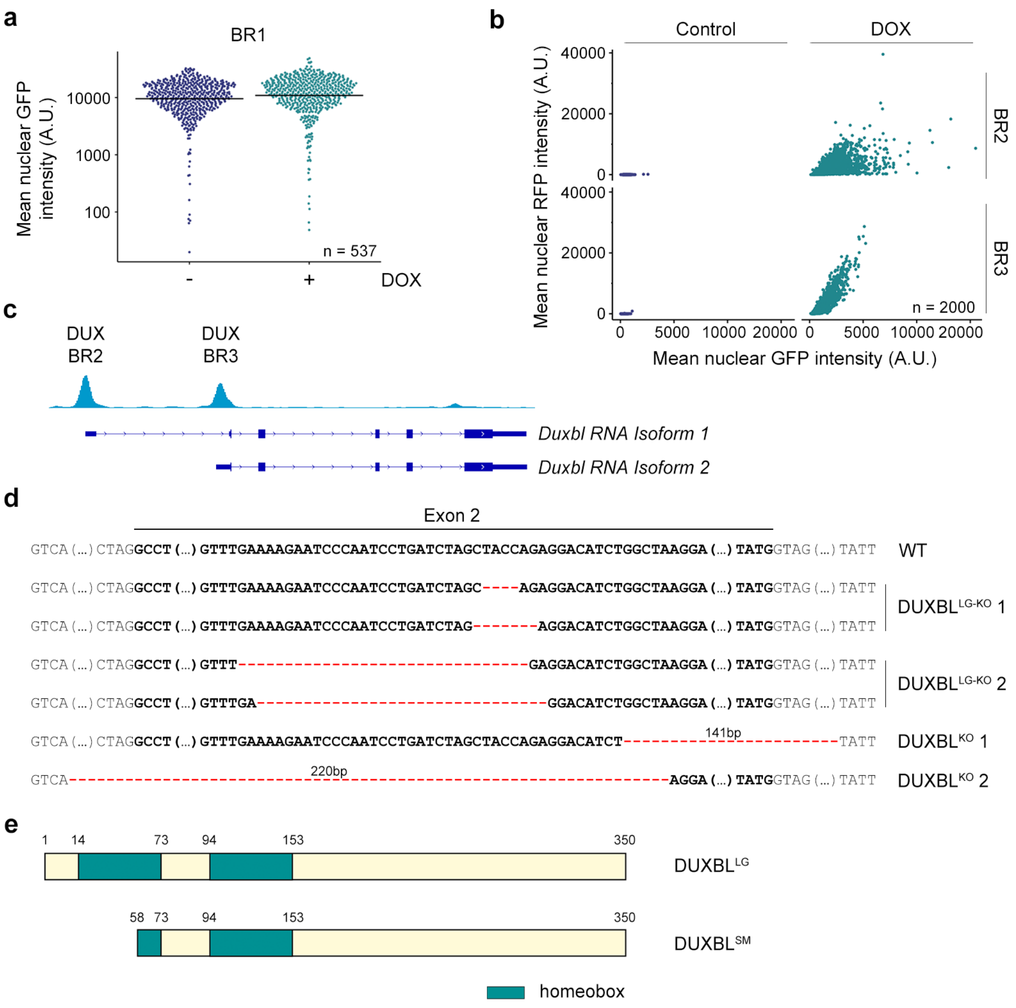
a
BR1
Mean nuclear GFP intensity (A.U.)
10000
1000
100
n = 537
-
+
DOX
b
Control
DOX
Mean nuclear RFP intensity (A.U.)
40000
20000
0
BR2
BR3
n = 2000
0
5000
10000
20000
Mean nuclear GFP intensity (A.U.)
c
DUX BR2
DUX BR3
Duxbl RNA Isoform 1
Duxbl RNA Isoform 2
d
Exon 2
GTCA (...) CTAGGCCT (...) GTTTGAAAAGAATCCCAATCCTGATCTAGCTACCAGAGGACATCTGGCTAAGGA (...) TATGGTAG (...) TATT
WT
GTCA (...) CTAGGCCT (...) GTTTGAAAAGAATCCCAATCCTGATCTAGC----AGAGGACATCTGGCTAAGGA (...) TATGGTAG (...) TATT
GTCA (...) CTAGGCCT (...) GTTTGAAAAGAATCCCAATCCTGATCTAG-------AGGACATCTGGCTAAGGA (...) TATGGTAG (...) TATT
DUXBL LG-KO 1
GTCA (...) CTAGGCCT (...) GTTT-----GAGGACATCTGGCTAAGGA (...) TATGGTAG (...) TATT
GTCA (...) CTAGGCCT (...) GTTTGA-----GGACATCTGGCTAAGGA (...) TATGGTAG (...) TATT
DUXBL LG-KO 2
GTCA (...) CTAGGCCT (...) GTTTGAAAAGAATCCCAATCCTGATCTAGCTACCAGAGGACATCT-----141bp-----TATT
DUXBL KO 1
GTCA-----220bp-----AGGA (...) TATGGTAG (...) TATT
DUXBL KO 2
e
1
14
73
94
153
350
DUXBL LG
58
73
94
153
350
DUXBL SM
homeobox

**Figure S1**

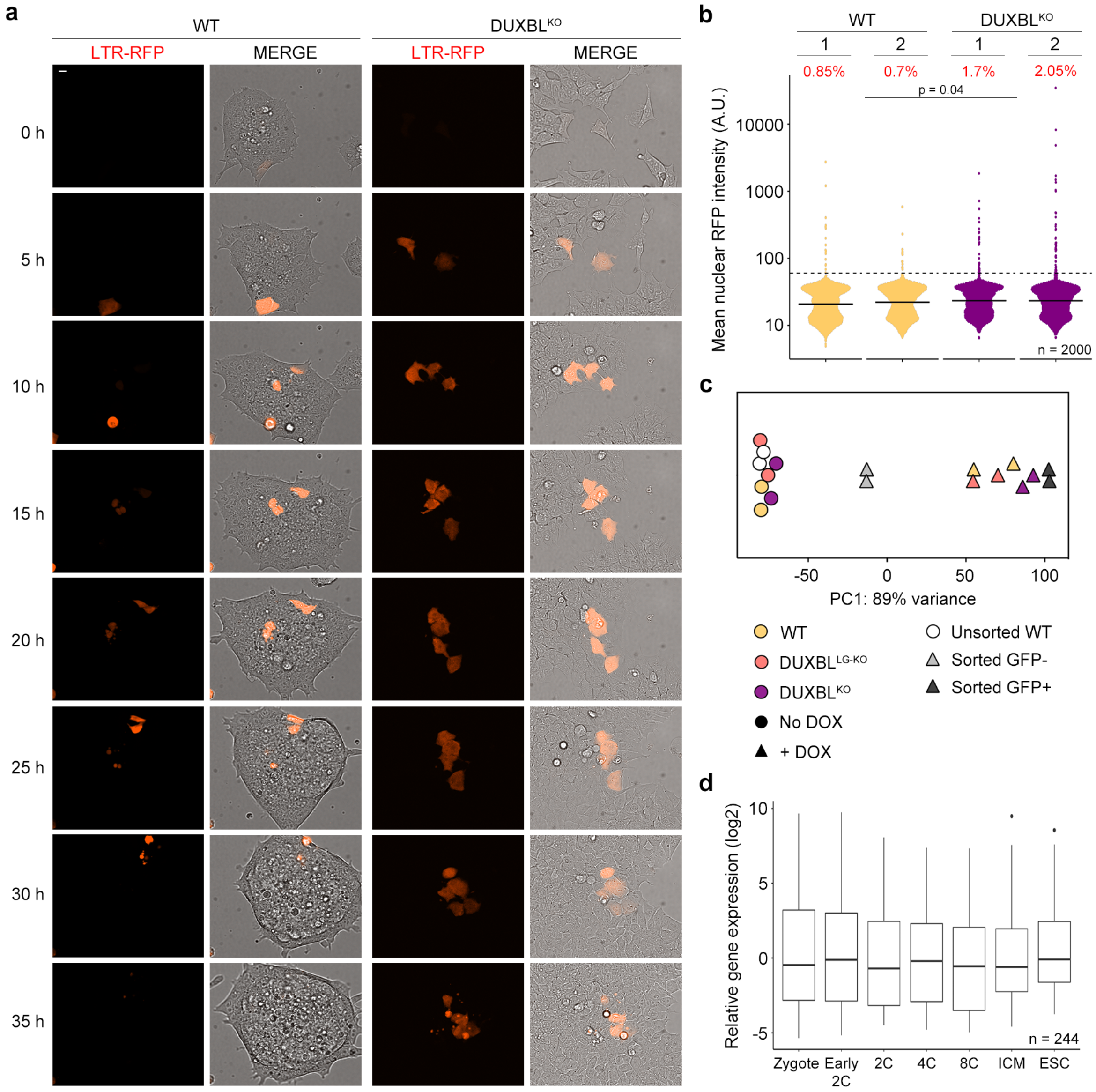

a
WT
DUXBL^KO
LTR-RFP
MERGE
LTR-RFP
MERGE
0 h
5 h
10 h
15 h
20 h
25 h
30 h
35 h
b
WT
DUXBL^KO
1
2
1
2
0.85%
0.7%
1.7%
2.05%
p = 0.04
Mean nuclear RFP intensity (A.U.)
10000
1000
100
10
n = 2000
c
-50
0
50
100
PC1: 89% variance
WT
DUXBL^LG-KO
DUXBL^KO
No DOX
+ DOX
Unsorted WT
Sorted GFP-
Sorted GFP+
d
Relative gene expression (log2)
10
5
0
-5
n = 244
Zygote
Early 2C
2C
4C
8C
ICM
ESC


**Figure S2**

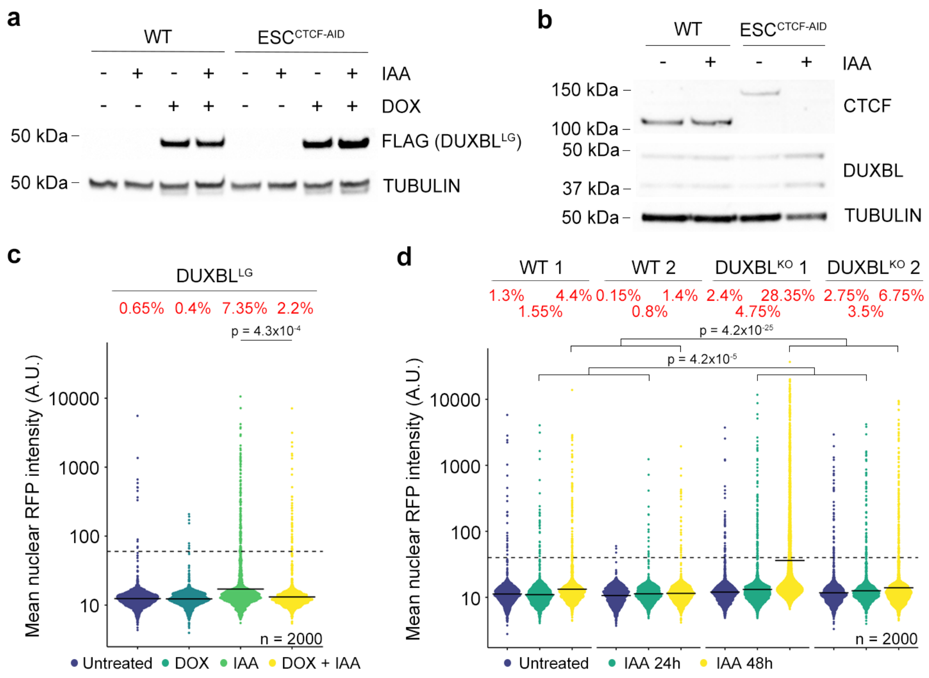
a
WT
ESC CTCF-AID
IAA
DOX
50 kDa
FLAG (DUXBL LG)
50 kDa
TUBULIN
b
WT
ESC CTCF-AID
IAA
150 kDa
100 kDa
CTCF
50 kDa
37 kDa
DUXBL
50 kDa
TUBULIN
c
DUXBL LG
0.65% 0.4% 7.35% 2.2%
p = 4.3x10-4
Mean nuclear RFP intensity (A.U.)
10000
1000
100
10
n = 2000
Untreated DOX IAA DOX + IAA
d
WT 1
WT 2
DUXBL KO 1
DUXBL KO 2
1.3% 4.4% 0.15% 1.4% 2.4% 28.35% 2.75% 6.75%
1.55% 0.8% 4.75% 3.5%
p = 4.2x10-25
p = 4.2x10-5
Mean nuclear RFP intensity (A.U.)
10000
1000
100
10
n = 2000
Untreated IAA 24h IAA 48h

**Figure S3**

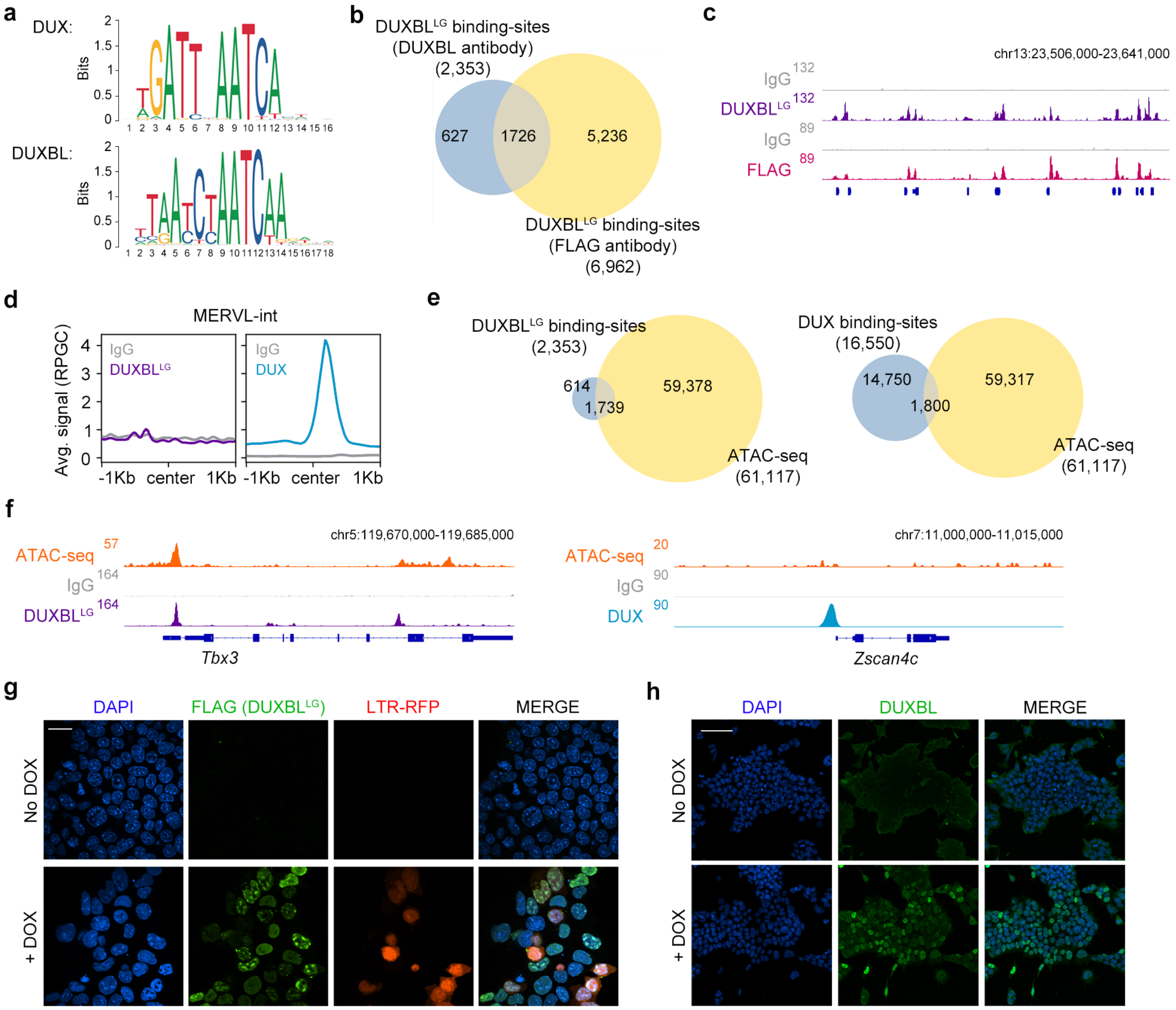
a
DUX:
DUXBL:
Bits
b
DUXBLLG binding-sites (DUXBL antibody) (2,353)
627
1726
5,236
DUXBLLG binding-sites (FLAG antibody) (6,962)
c
chr13:23,506,000-23,641,000
IgG
DUXBLLG
FLAG
d
MERVL-int
Avg. signal (RPGC)
-1Kb center 1Kb
DUX
e
DUXBLLG binding-sites (2,353)
614
1,739
59,378
ATAC-seq (61,117)
DUX binding-sites (16,550)
14,750
1,800
59,317
f
chr5:119,670,000-119,685,000
chr7:11,000,000-11,015,000
Tbx3
Zscan4c
g
DAPI
FLAG (DUXBLLG)
LTR-RFP
MERGE
No DOX
+ DOX
h
DUXBL

Figure S4

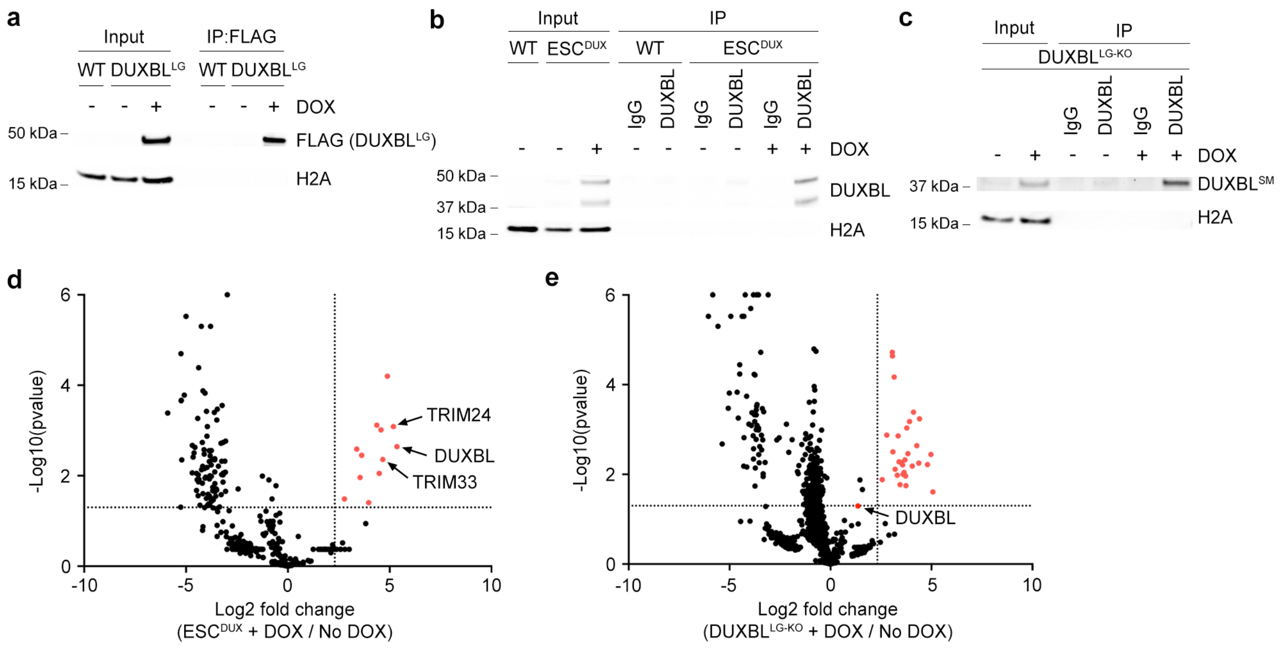
a
Input
IP:FLAG
WT DUXBLLG
WT DUXBLLG
DOX
50 kDa
15 kDa
FLAG (DUXBLLG)
H2A
b
Input
IP
WT ESCDUX
WT
ESCDUX
IgG
DUXBL
DOX
50 kDa
37 kDa
15 kDa
DUXBL
H2A
c
Input
IP
DUXBLLG-KO
IgG
DUXBL
DOX
37 kDa
15 kDa
DUXBLSM
H2A
d
-Log10(pvalue)
TRIM24
DUXBL
TRIM33
Log2 fold change
(ESCDUX + DOX / No DOX)
e
-Log10(pvalue)
DUXBL
Log2 fold change
(DUXBLLG-KO + DOX / No DOX)

**Figure S5**

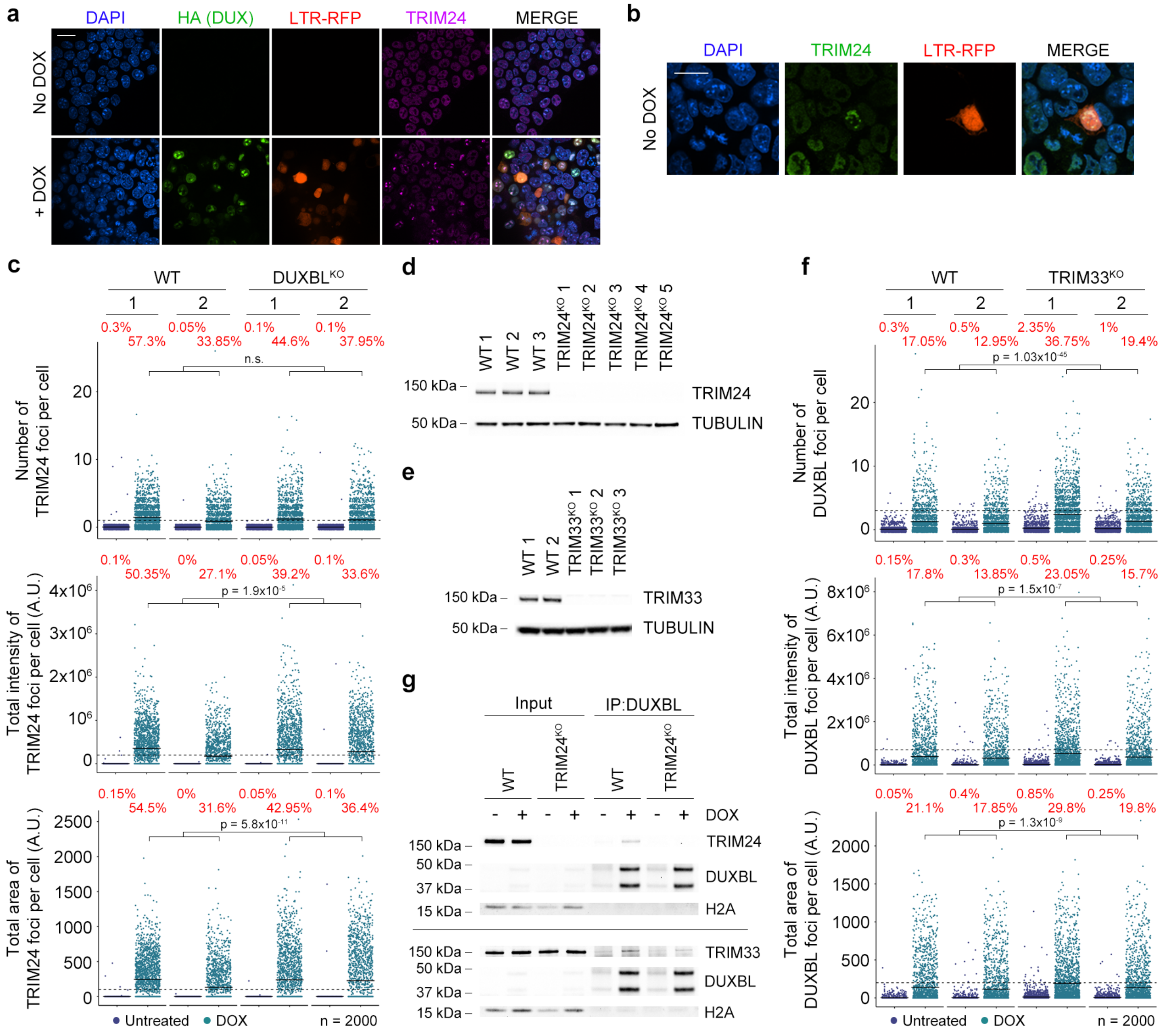

a
DAPI
HA (DUX)
LTR-RFP
TRIM24
MERGE
No DOX
+ DOX
b
DAPI
TRIM24
LTR-RFP
MERGE
No DOX
c
WT
DUXBL^KO
Number of TRIM24 foci per cell
Total intensity of TRIM24 foci per cell (A.U.)
Total area of TRIM24 foci per cell (A.U.)
n.s.
p = 1.9x10^-5
p = 5.8x10^-11
Untreated
DOX
n = 2000
d
WT 1
WT 2
WT 3
TRIM24^KO 1
TRIM24^KO 2
TRIM24^KO 3
TRIM24^KO 4
TRIM24^KO 5
150 kDa
50 kDa
TRIM24
TUBULIN
e
TRIM33^KO 1
TRIM33^KO 2
TRIM33^KO 3
TRIM33
g
Input
IP:DUXBL
DOX
37 kDa
15 kDa
DUXBL
H2A
f
TRIM33^KO
Number of DUXBL foci per cell
Total intensity of DUXBL foci per cell (A.U.)
Total area of DUXBL foci per cell (A.U.)
p = 1.03x10^-45
p = 1.5x10^-7
p = 1.3x10^-9


**Figure S6**

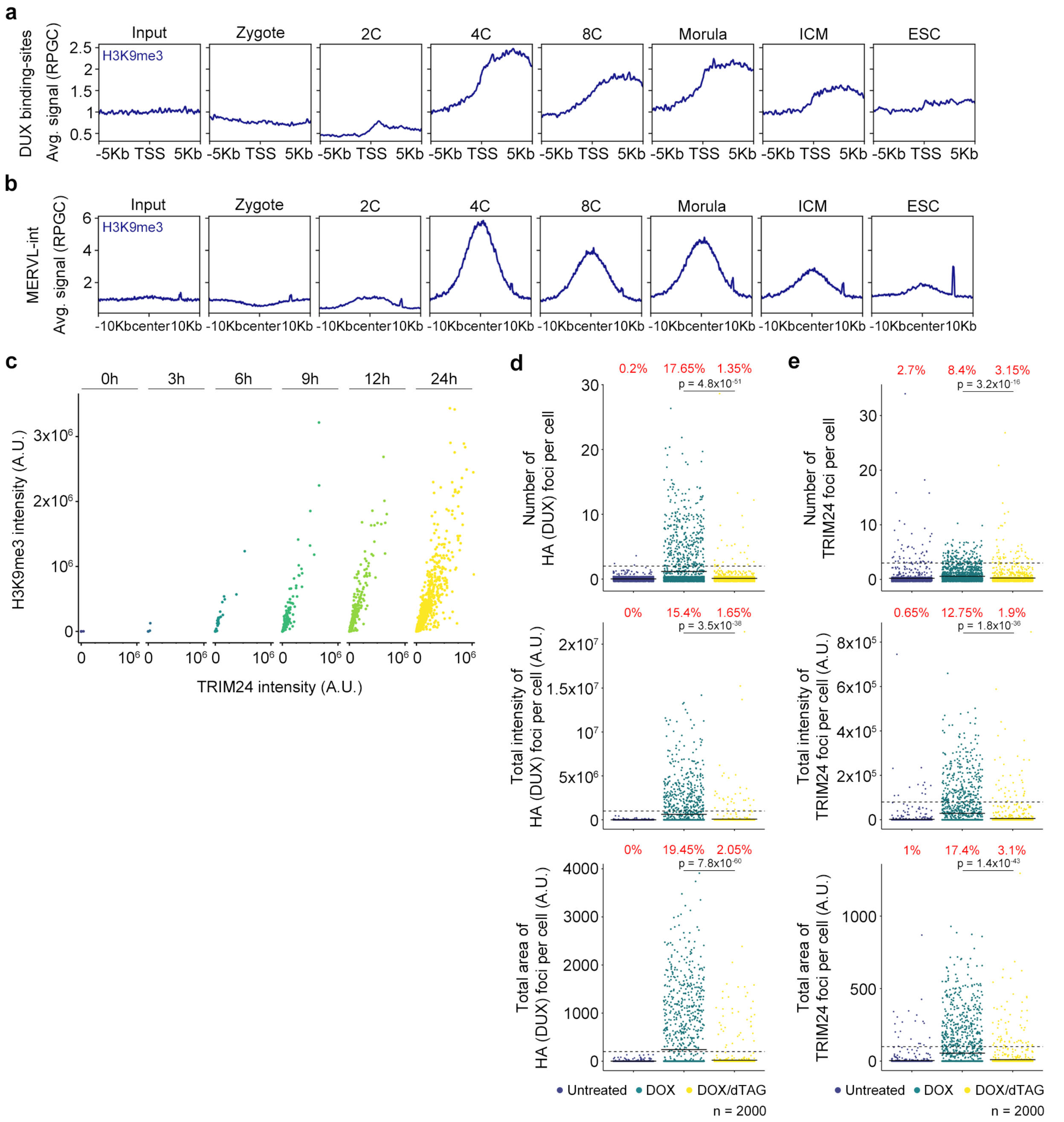

a
Input
Zygote
2C
4C
8C
Morula
ICM
ESC
H3K9me3
DUX binding-sites
Avg. signal (RPGC)
-5Kb TSS 5Kb
b
MERVL-int
Avg. signal (RPGC)
-10Kb center 10Kb
c
0h
3h
6h
9h
12h
24h
H3K9me3 intensity (A.U.)
TRIM24 intensity (A.U.)
d
0.2% 17.65% 1.35%
p = 4.8x10-51
Number of HA (DUX) foci per cell
0% 15.4% 1.65%
p = 3.5x10-38
Total intensity of HA (DUX) foci per cell (A.U.)
0% 19.45% 2.05%
p = 7.8x10-60
Total area of HA (DUX) foci per cell (A.U.)
Untreated DOX DOX/dTAG
n = 2000
e
2.7% 8.4% 3.15%
p = 3.2x10-16
Number of TRIM24 foci per cell
0.65% 12.75% 1.9%
p = 1.8x10-36
Total intensity of TRIM24 foci per cell (A.U.)
1% 17.4% 3.1%
p = 1.4x10-43
Total area of TRIM24 foci per cell (A.U.)
Untreated DOX DOX/dTAG
n = 2000


### Figure S7

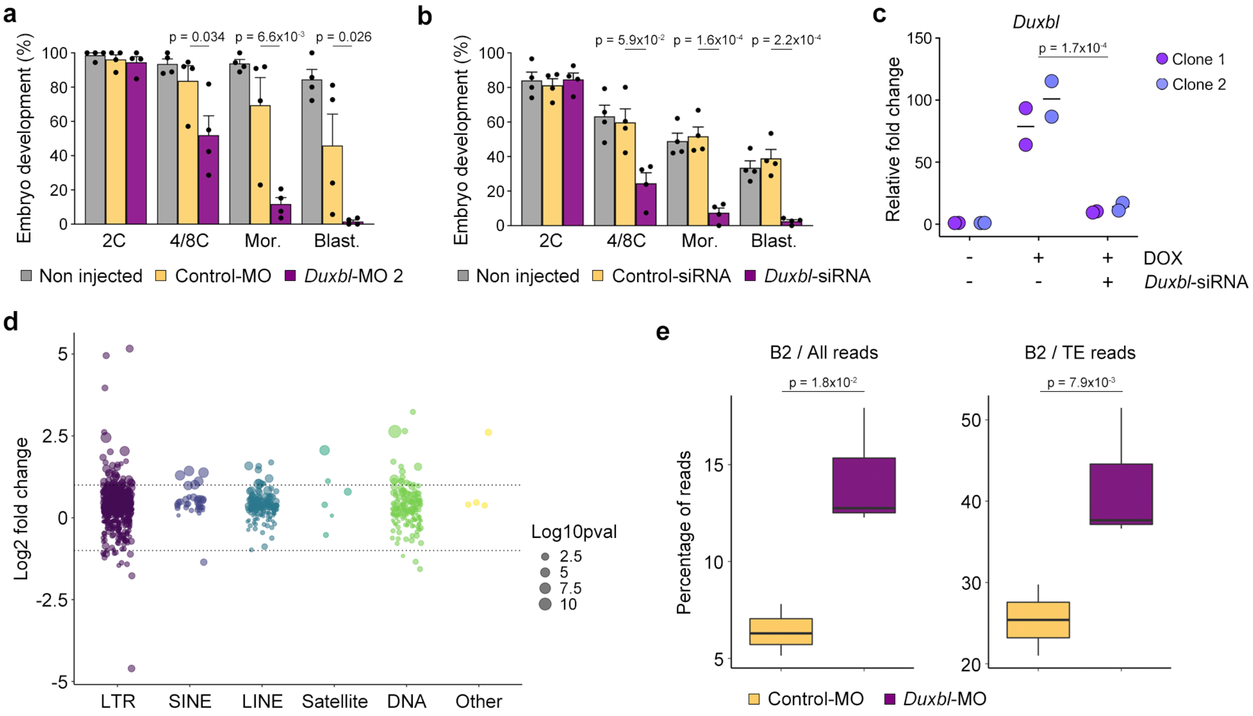
a
Embryo development (%)
p = 0.034
p = 6.6x10-3
p = 0.026
2C
4/8C
Mor.
Blast.
Non injected
Control-MO
Duxbl-MO 2
b
p = 5.9x10-2
p = 1.6x10-4
p = 2.2x10-4
Non injected
Control-siRNA
Duxbl-siRNA
c
Duxbl
Relative fold change
p = 1.7x10-4
Clone 1
Clone 2
DOX
Duxbl-siRNA
d
Log2 fold change
Log10pval
LTR
SINE
LINE
Satellite
DNA
Other
e
B2 / All reads
B2 / TE reads
p = 1.8x10-2
p = 7.9x10-3
Percentage of reads
Control-MO
Duxbl-MO

**Figure S8**

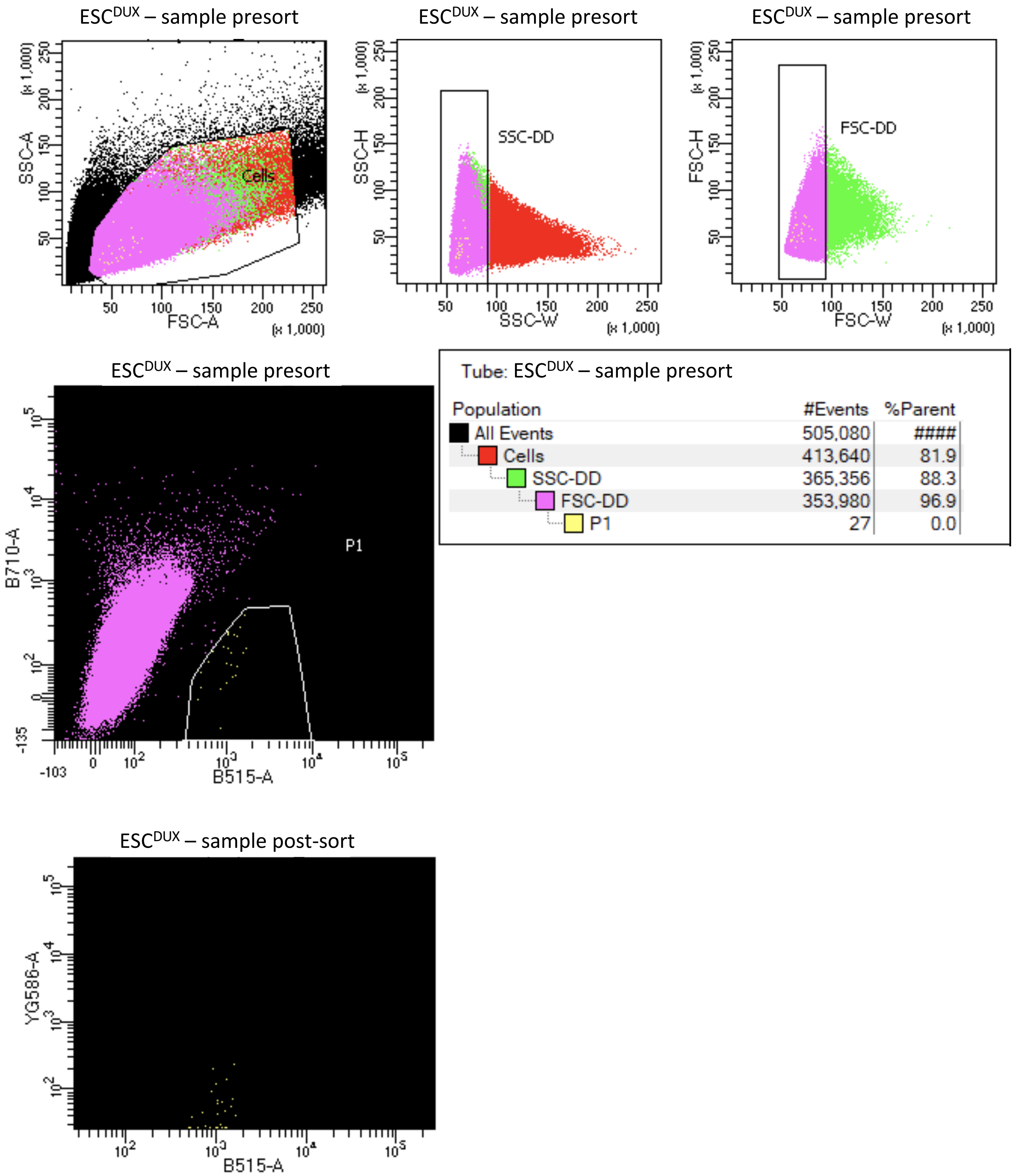


Tube: ESC$^{DUX}$ – sample presort

| Population | #Events | %Parent |
|---|---|---|
| All Events | 505,080 | #### |
| Cells | 413,640 | 81.9 |
| SSC-DD | 365,356 | 88.3 |
| FSC-DD | 353,980 | 96.9 |
| P1 | 27 | 0.0 |

**Figure S9**
